# Rap1b Activates Endosomal AC9 to Drive the Second cAMP Wave

**DOI:** 10.64898/2026.05.06.723328

**Authors:** Xuefeng Zhang, Alejandro Pizzoni, Alipio Pinto, Daniel L. Altschuler

## Abstract

GsPCR signaling orchestrates cAMP production in three distinct spatial waves originating from the plasma membrane (PM), endosomes, and the nucleus. While the molecular drivers of the PM and nuclear waves are well defined, the regulation of the endosomal cAMP wave remains insufficiently understood. Here, we identify the small GTPase Rap1b as a direct activator of the endosomal adenylyl cyclase 9 (AC9), revealing a novel mechanism for intracellular cAMP synthesis. Like Gαs, Rap1b-GTP interacts with the C2 domain of AC9, allosterically enhancing its catalytic activity both *in vitro* and in cells. Using AC9 mutations that selectively disrupt Rap1b versus Gαs binding, we elucidate the role of the Rap1b–AC9 unit in mediating the endosomal cAMP wave, introducing a new layer of spatial regulation to cAMP signaling.

## Introduction

G protein–coupled receptors (GPCRs), the largest family of mammalian membrane receptors, control diverse physiological processes, thus representing major drug targets (*1, 2*). Ligand binding induces conformational changes that activate heterotrimeric G proteins (Gαβγ), driving GDP–GTP exchange on Gα, dissociation of Gα–GTP from Gβγ, and regulation of downstream effectors by both subunits. Within this family, Gs–coupled GPCRs (GsPCRs) activate adenylyl cyclases (ACs), generating the second messenger cyclic AMP (cAMP). The mammalian genome encodes ten AC isoforms (AC1–AC10). While AC10 (soluble adenylyl cyclase; sAC) is G–protein–independent, AC1–AC9 are positively regulated by Gαs–GTP and membrane–bound (*3*).

AC1–AC9 isoforms share a conserved architecture: two six–helix transmembrane bundles (TM1–6 and TM7–12) connected to a cytosolic core via two helical domains (*4*). The cytosolic core includes pseudo–homologous catalytic domains C1a and C2a flanked by non–homologous regulatory modules C1b and C2b. The C1a–C2a dimer forms adjacent catalytic and allosteric (forskolin–binding) pockets, and recent structural analyses show that Gαs engages ACs primarily via its switch II helix. This helix inserts into a cleft of C2a and induces conformational changes in C1a and C2a, including in a forskolin–binding helix of C2a, thereby optimizing catalytic alignment and allosteric coupling (*5, 6*).

Historically, cAMP signaling was thought to be confined to the PM. Subsequent work has shown that ligand-activated GsPCRs generate a ‘first’ cAMP wave at the PM and, after endocytosis, a ‘second’ endosomal wave that drives nuclear responses (*7, 8*). Our recent work further proposes a ‘third’ cAMP wave generated by Ca²⁺-dependent activation of nuclear sAC (*9*). Despite the high diffusivity of cAMP (*10*), its effective range in cells appears limited to ∼30–60 nm (*11–17*), implying that coupling between waves relies on local signaling modules. How cAMP signals are integrated across these compartments—and what additional factors link them—remains unclear. In our three-wave model, the first wave is generated by canonical Gαs–AC activity at the PM and the third by the Ca²⁺-dependent nuclear sAC. However, the identity of the AC isoform driving the endosomal second wave, as well as the machinery that supports its activity, is still unknown (*18*).

In previous work, we showed that CAP1 scaffolds AC and Rap1b into a functional unit that modulates cAMP dynamics in mammalian cells: CAP1’s C–terminal β–barrel binds Rap1b in a GTP–independent manner, while its N–terminus engages the C1a domain of AC (*19, 20*). This arrangement leaves Rap1b’s effector domains accessible and likely positions Rap1b at the AC catalytic interface, providing a scaffold for non–canonical activation of cAMP synthesis. Here, we identify AC9 as the sole isoform in this pathway and show that Rap1b regulates AC activity directly and independently of Gαs: Rap1b–GTP binds AC9 and inserts its switch II helix into a cleft of C2a, thereby activating AC9 and markedly potentiating the otherwise weak effect of forskolin on this isoform (*6*). Guided by computational modeling and the AC9–Gαs cryo–EM structure, we pinpointed AC9 mutations that differentially control Gαs– versus Rap1b–mediated activation. Collectively, these results demonstrate that Rap1b can trigger endosomal AC9 activity independently of Gαs to generate the second cAMP wave.

## Results

### AC9 is the only CAP1–sensitive isoform in intact cells

We recently reported that CAP1 binds and activates mammalian AC *in vitro* and in living cells (*20*). To assess the isoform specificity of the CAP1–AC regulation in cells, we used the HC–1 cell line, which, as reported (*21*), endogenously express β2–AR and Gs but do not respond to isoproterenol (ISO) or forskolin (FK) unless an AC is transfected (*22*), making them an ideal model to test CAP1 effects on specific ACs without an endogenous AC background. Based on their properties, AC1–AC9 form four groups (*23*). We expressed one AC isoform representative of each group—AC3 (group I), AC2 (group II), AC5 (group III), and AC9 (group IV)—and either overexpressed CAP1 or knocked it down with sh–CAP1, then quantified ISO–stimulated cAMP dynamics using the cytosolic FRET sensor H188 (*24*). All tested isoforms were functional and restored the ISO response, but only AC9 was modulated by CAP1 (Figs. 1A, S1A, and S1C). Thus, despite the lack of isoform specificity *in vitro* with purified components (*20*), AC9 is the only CAP1–sensitive isoform in intact living cells.

**Fig. 1.**
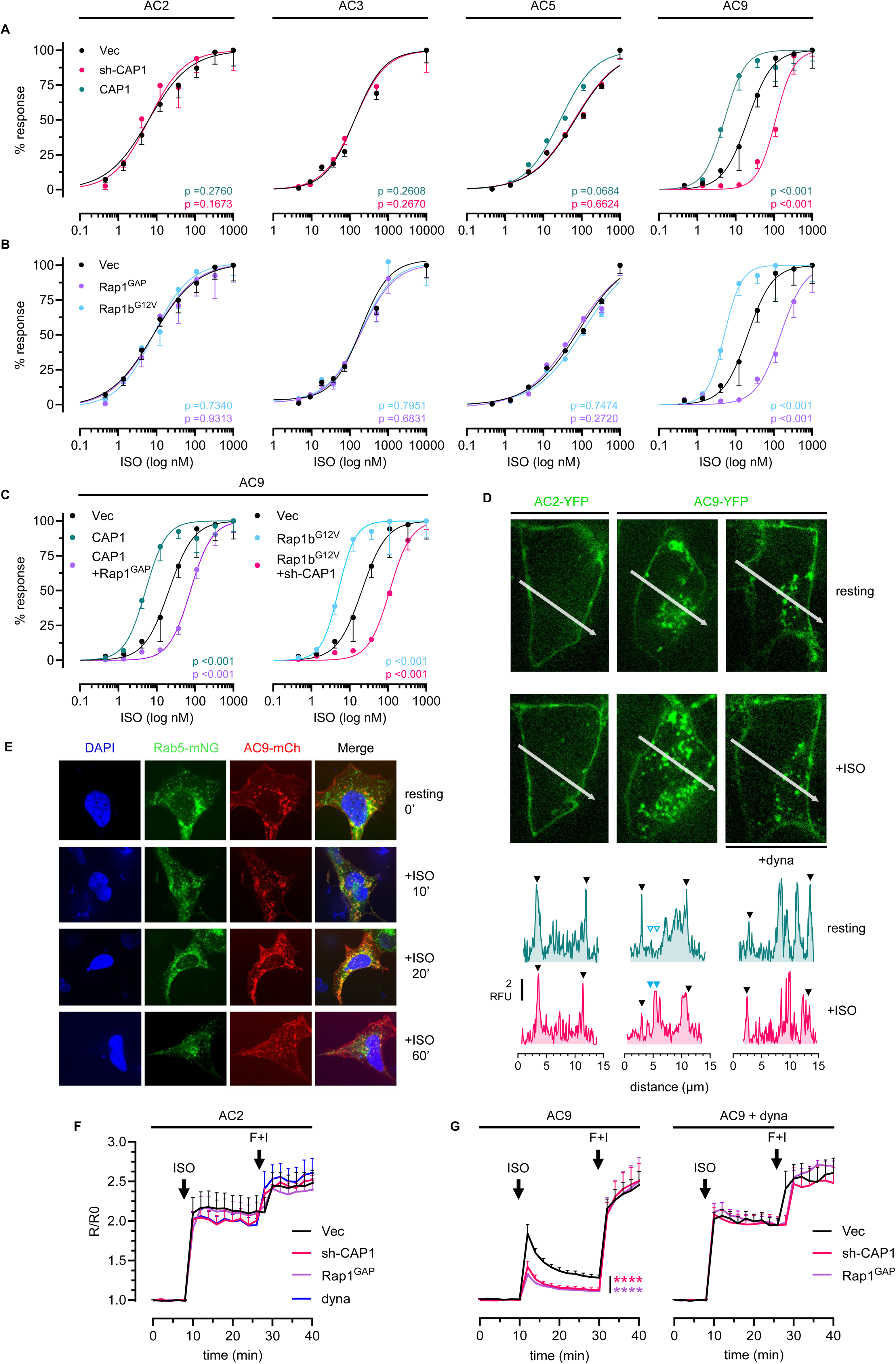
CAP1–Rap1b specifically regulates AC9 signaling by modulating its internalization–dependent second cAMP wave. Dose–response (DR) curves show normalized responses (% max) calculated from ΔR/R₀ Förster resonance energy transfer (FRET) ratios recorded with the H188 cAMP sensor 20 min after the addition of isoproterenol (ISO). **(A)** Effect of CAP1 overexpression (CAP1) or knockdown (sh–CAP1) on cAMP production in HC–1 cells transfected with AC2, AC3, AC5, or AC9 and stimulated with ISO in a DR protocol. **(B)** Effect of Rap1b activation by the constitutively active mutant Rap1b^G12V^ or Rap1b inactivation by the GTPase–activating protein Rap1^GAP^ on cAMP production in the same cells under ISO DR conditions. AC3–expressing cells were pretreated with 150μM carbachol (CCH). **(C)** Combined effect of CAP1 and Rap1b modulation in ISO DR experiments: CAP1 overexpression together with Rap1^GAP^ (left) and sh–CAP1 together with constitutively active Rap1b^G12V^ (right). **(D)** Representative frames from live–cell confocal time–lapse imaging showing the subcellular localization of AC2–YFP (left panels) and AC9–YFP (middle and right panels) before stimulation (resting) and after ISO (30nM) in the same HC–1 cells. Dynasore (dyna; 70μM) was used to block endocytosis. Bottom panels: quantification of YFP fluorescence along line scans across the entire cell at rest and 20 min after ISO. Black arrows indicate PM, whereas blue arrows indicate AC9–YFP internalization. **(E)** Representative confocal micrographs from fixed HC–1 cells showing colocalization of Rab5–mNG and AC9–mCherry at rest and 10, 20, or 60 min after ISO (30nM) in different cells. **(F, G)** Real–time cytosolic cAMP measurements in HC–1 cells expressing AC2 (F) or AC9 (G) and stimulated with ISO (30nM), followed by forskolin and 3–isobutyl–1–methylxanthine (IBMX) (F+I; FK 20μM, IBMX 250μM), using the H188 sensor. Traces represent normalized FRET ratios (R/R₀). Dynasore (dyna; 70μM) was used to block endocytosis. DR data (panels A, B, and C; pooled from ≥3 independent experiments) were fit by nonlinear regression to a four–parameter logistic equation to obtain EC₅₀ values with 95% confidence intervals (CI). Differences (vs. vector) were assessed by extra–sum–of–square; p-values reported in Fig. S2A. Traces (panels F and G) are shown as mean ± SEM; n ≥ 7 cells from ≥ 3 independent experiments. Significance was assessed at t = 20 min after ISO (vertical line) by one-way ANOVA with Dunnett’s multiple-comparisons test versus vector control. Color-matched asterisks denote significance (no asterisks, not significant): ****p < 0.0001.

### AC9 is the only Rap1b–GTP–sensitive AC isoform in intact cells

Given our prior finding that CAP1 binds prenylated Rap1b (*19*), we asked whether CAP1 regulation of AC9 depends on Rap1b. We co–expressed AC isoforms with Rap1^GAP^ or Rap1b^G12V^ in HC–1 cells and observed, as seen with CAP1, Rap1b–GTP sensitivity only for AC9 (Figs. 1B and S1B–C). When CAP1 was downregulated, Rap1b^G12V^ no longer enhanced signaling, and when Rap1b was inactivated by Rap1^GAP^, overexpressed CAP1 also lost its potentiating effect (Figs. 1C and S1D–E). Endogenous sAC, assayed by its bicarbonate response, was unaffected by CAP1–Rap1b, ruling out sAC as the source of these differences (Fig. S1F). These combined results indicate that the effects of CAP1 and Rap1b on AC9 are interdependent, thus requiring a working CAP1–Rap1b complex.

### CAP1–Rap1b modulates AC9 after internalization

AC9 internalizes upon ISO stimulation (*25*), whereas our preliminary data indicated that AC2 does not. In unstimulated HC–1 cells, both isoforms localized predominantly to the PM; only AC9 exhibited, in addition, an intracellular perinuclear pool (Fig. 1D). Upon ISO stimulation, only AC9 internalized via endocytosis (Fig. 1D), into a Rab5^+^compartment (Fig. 1E). To define how these different AC trafficking profiles shape cAMP signaling dynamics, we compared AC9 and AC2. Non–internalized AC2 generated only a sustained first cAMP wave (*i.e.*, PM) that was fully insensitive to CAP1–Rap1b (Fig. 1F). By contrast, for AC9, ISO triggered an internalization–dependent second cAMP wave that was reduced by CAP1–Rap1b downregulation (Fig. 1G). CAP1–Rap1b manipulation did not affect ISO–mediated AC9 internalization (Fig. S1G). Thus, these combined results indicate a post-internalization effect of CAP1–Rap1b on AC9-mediated cAMP synthesis.

### Rap1b directly engages AC9’s C2 domain

As described above, CAP1–Rap1b controls specifically AC9 over AC2. Aiming to find the specific AC9 domain involved, we built chimeras of AC9 (CAP1–Rap1b sensitive) and AC2 (CAP1–Rap1b insensitive) (Fig. S2A). While most chimeras were nonfunctional, AC9(1037)AC2, an AC9 backbone carrying the AC2–C2 domain, restored the ISO–evoked response in HC–1 cells (Fig. S2B). This chimera (Fig. 2A) internalized upon stimulation (Fig. 2B) but completely lost CAP1–Rap1b sensitivity (Figs. 2C–D and S2C), suggesting the involvement of the AC9–C2 domain. To test whether Rap1b binds directly to AC9 C2, we used microscale thermophoresis (MST) with AC9–YFP as the probe, and Rap1b–GTP and Gαs–GTP as titrants. MST confirmed a direct AC9–YFP::Rap1b–GTP binding (Kd ∼ 1.3 µM), modestly weaker than AC9–YFP::Gαs–GTP (Kd ∼ 0.6 µM) (Fig. 2E). Pull–downs with purified His–C1a and His–C2a against HA–Rap1b–GTP, revealed a specific Rap1b–GTP::C2a interaction (Fig. 2F). MST assays using GFP–Rap1b^G12V^ as probe, confirmed the interaction with the C2a domain (Fig. 2G, Kd ∼ 9 µM). Thus, the combined results indicate that Rap1b binds directly to the AC9–C2a domain (Fig. 2H).

**Fig. 2.**
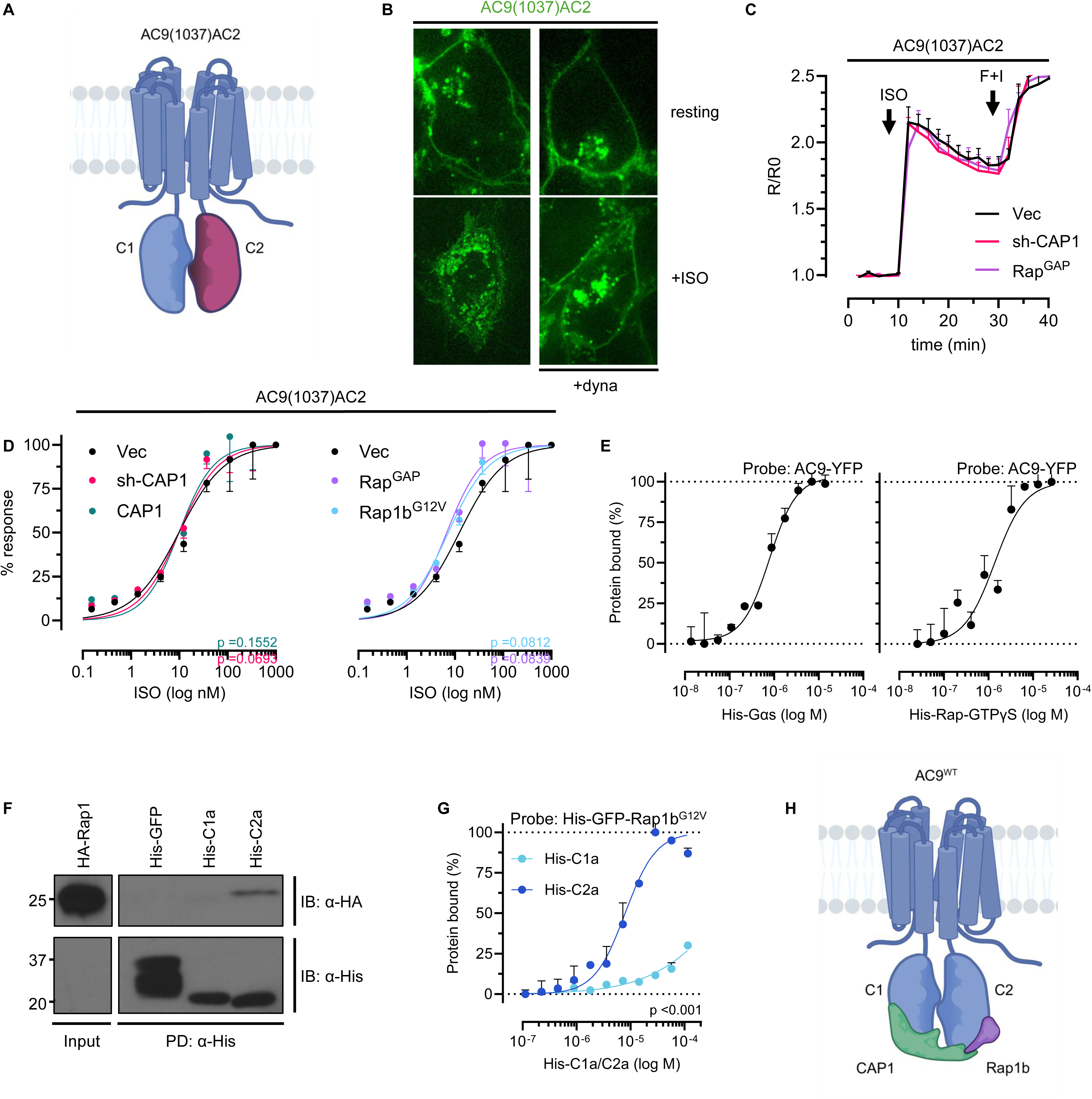
AC9’s C2 domain is directly engaged by Rap1b, enabling isoform–specific regulation. **(A)** Schematic representation of the AC9(1037)AC2 chimera, an AC9 backbone carrying the AC2 C2 domain. **(B)** Representative confocal immunofluorescence micrographs (anti–HA) showing the subcellular localization of the HA–tagged AC9(1037)AC2 chimera at rest and after ISO (30nM) stimulation. Dynasore (dyna; 70μM) was used to block endocytosis–related morphological changes. **(C)** Real–time cytosolic cAMP measurements in HC–1 cells expressing the AC9(1037)AC2 chimera and the H188 FRET sensor during ISO (30nM) stimulation followed by forskolin and IBMX (F+I; FK 20μM, IBMX 250μM). Traces represent normalized FRET ratios (R/R₀). **(D)** Dose–response (DR) effects of CAP1 overexpression (CAP1) or knockdown (sh–CAP1) (left) and Rap1b modulation by constitutively active Rap1b^G12V^ or the GTPase–activating protein Rap1^GAP^ (right) on cAMP production in HC–1 cells expressing the AC9(1037)AC2 chimera stimulated with ISO (30nM). ΔR/R₀ (%) FRET responses were measured with the H188 sensor. **(E)** Microscale thermophoresis (MST) analysis of AC9–YFP binding to purified His–Gαs (left) or His–Rap1b–GTPγS (right) *in vitro*. **(F)** Representative pull–down assay showing interaction between Rap1b and the AC9 C2a domain. HEK293 cells were transfected with HA–Rap1b^G12V^, and lysates were incubated with Ni–NTA agarose beads loaded with His–tagged AC9 C2a. Complex formation was detected by immunoblotting with an HA–specific antibody. Representative experiment (n = 3). **(G)** MST analysis of His–GFP–Rap1b^G12V^ binding to purified His–C1a or His–C2a domains *in vitro*. **(H)** Schematic representation of CAP1–Rap1b binding to the AC9 C2 domain. Traces (panel C) are shown as mean ± SEM; n ≥ 8 cells from ≥ 3 independent experiments. Significance was assessed at t = 20 min after ISO (vertical line) by one-way ANOVA with Dunnett’s multiple-comparisons test versus vector control. Absence of asterisks denotes non-significance. DR data (panels D, E, G); pooled from n=3 independent experiments) were fit by nonlinear regression to a four–parameter logistic equation to obtain EC₅₀ values with 95% confidence intervals (CI). Differences (vs. vector) were assessed by extra–sum–of–square; p-values reported in Fig. S2E (panel D). EC₅₀ values were 6.0 × 10⁻⁷ M for AC9::Gαs and 1.27 × 10⁻⁶ M for AC9::Rap1b (p < 0.0001) (panel E). EC₅₀ values were 6.56 × 10⁻⁵ M for C1a and 9.66 × 10⁻⁷ M for C2a (p < 0.0001) (panel G).

### Rap1b–GTP activates AC9 *in vitro*

The convergence of Gαs and Rap1b–GTP binding on C2 prompted us to test whether Rap1b–GTP competes with Gαs–GTP at this site. Purified Rap1b–GTP dose–dependently reduced Gαs–GTP–mediated AC9 activation (Fig. 3A, left), consistent with a competitive displacement. However, inhibition remained partial even at saturating Rap1b–GTP, raising the question of whether Rap1b–GTP can directly activate AC9 without Gαs. Indeed, purified Rap1b–GTP activated AC9, albeit with lower efficacy than Gαs–GTP (Fig. 3A, right).

**Fig. 3.**
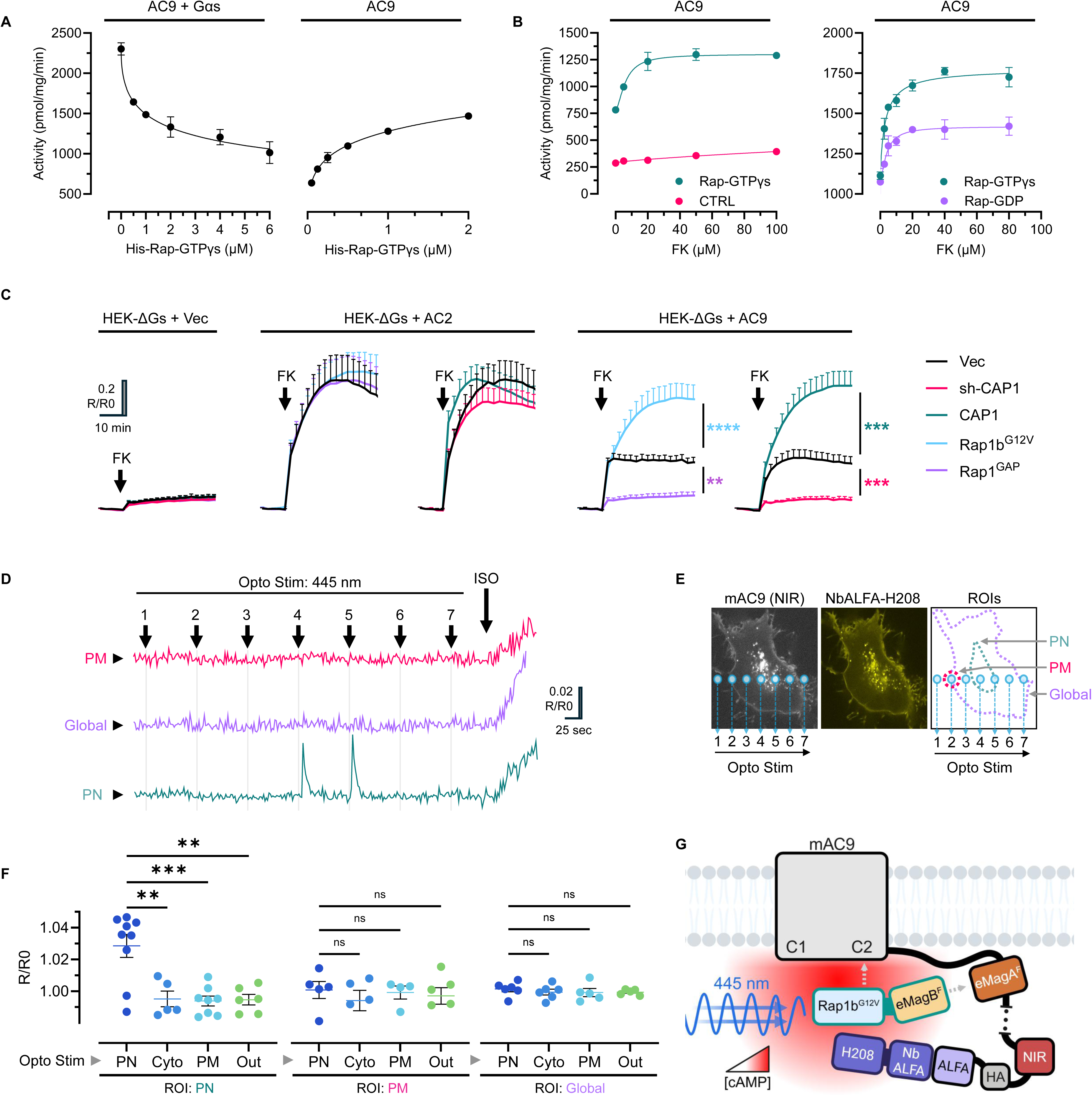
Rap1b–GTP directly activates AC9. **(A)** Dose–dependent effect of purified Rap1b–GTP on Gαs–GTP–mediated AC9 activation *in vitro*. **(B)** Rap1b–GTP potentiates forskolin (FK)–induced AC9 activity in a GTP–dependent manner, consistent with allosteric coupling between the Rap1b–binding site and the FK–binding site. **(C)** Real–time cAMP measurements using the H188 Förster resonance energy transfer (FRET) cAMP sensor in HEK–ΔGαs cells. Traces represent normalized FRET ratios (R/R₀). Stimulation with FK (75 μM) alone elicited minimal responses (left). Transfection with AC2 (middle) or AC9 (right) restored FK sensitivity. AC2–mediated responses remained insensitive to CAP1 or Rap1b modulation, whereas AC9 activity was modulated by CAP1 overexpression or knockdown (sh–CAP1) and by constitutively active Rap1b (Rap1b^G12V^) or Rap1^GAP^–mediated Rap1b inactivation. **(D)** Real–time cAMP measurements using the NbALFA–H208 FRET sensor in HC–1 cells coexpressing mAC9 and Rap1b^G12V^–eMagB^F^ during optogenetic stimulation with blue light (445 nm). Representative experiment showing stimulation of mAC9 in different cell areas (Opto Stim; 1–7, shown in panel E) and measured using different ROIs (PM, plasma membrane; global; PN, perinuclear). **(E)** Confocal micrographs of the same HC–1 cell used for the traces in panel D. Images show NIR fluorescence from mAC9 (left) and YFP from NbALFA–H208 (middle). Optogenetically stimulated areas (Opto stim; 1–7) are indicated. **(F)** Quantification of the maximal ΔR/R₀ (%) FRET response during optogenetic stimulation in selected cytoplasmatic regions (Opto Stim; PN, Cyto, PM, Out) and measured in the indicated ROIs (PN, PM, Global). **(G)** Schematic representation of the optogenetic mAC9 experiment. Blue light (445 nm) recruits Rap1b^G12V^–eMagB^F^ to eMagAF–mAC9, promoting Rap1b^G12V^–mAC9 interaction and generating a local cAMP increase that is detected by NbALFA–H208 bound to the ALFA tag on mAC9. Traces in panel C are shown as mean ± SEM; n ≥ 8 cells from ≥ 3 independent experiments. Statistical significance was assessed at t = 20 min after FK addition (vertical line). The scatter plot in panel F (with mean ± SEM) represents individual cells. Data are from n = 3 independent experiments. Panels C and F: statistical significance versus vector control was assessed by one-way ANOVA with Dunnett’s multiple-comparisons test. Color-matched (or black) asterisks denote significance (ns/no asterisks: not significant); **p < 0.01, ***p < 0.001, ****p < 0.0001.

AC9 is insensitive to FK and requires Gαs for robust activity. (*6, 26*). Notably, Rap1b–GTP potentiated the effect of FK in a GTP–dependent manner (Fig. 3B). Thus, *in vitro*, Rap1–GTP recapitulates key features of Gαs–GTP regulation of AC9, activating the enzyme via allosteric coupling to the FK–binding site.

### Rap1b drives AC9 signaling independently of Gαs in live cells

To test whether Rap1b activates AC9 independently of Gαs, we used HEK293 cells lacking both Gαs and Gαolf (HEK–ΔGαs) (*27*), in which AC3 and AC6 are the predominant FK–sensitive isoforms (*28*). In this background, FK responses were small and CAP1–Rap1b–insensitive, consistent with FK–Gαs allostery (Fig. 3C). Overexpression of AC2 or AC9 greatly increased FK responsiveness, but, as in HC–1 cells, only AC9 conferred CAP1–Rap1b sensitivity (Fig. 3C). To test whether Rap1b–GTP alone is sufficient to activate AC9, we engineered a multi–tagged AC9 (mAC9) bearing an eMagA^F^ (*29*) domain for optogenetic recruitment of eMagB^F^–Rap1b^G12V^ and an ALFA tag (*30*) to tether a NbALFA–H208 cAMP sensor (*31*) for local cAMP readout (Fig. 3G and S3A). mAC9 was fully functional and retained localization (Fig. S3B–E). Blue–light recruitment of Rap1b^G12V^ to the intracellular mAC9 pool elicited a rapid, reversible local cAMP rise, whereas recruitment to other locations, including PM, was ineffective (Fig. 3D–G). Control experiments without eMagB^F^–Rap1b^G12V^ are shown in Fig. S3F. Thus, even in the absence of Gαs, CAP1–Rap1b is sufficient to activate specifically intracellular AC9.

### Rap1b switch II engages the AC9 C2a cleft

Next, we modeled the AC9–Rap1b complex with AlphaFold–Multimer (*32*), using the AC9 C1–C2 catalytic core and Rap1b^G12V^ (1–166) sequences. The top–ranked model positions Rap1b’s switch II (α2 helix and α2–β3 loop) into the cleft of AC9 C2a, mirroring the placement of the Gαs switch II helix seen in the AC9–Gαs cryo–EM structure (*5, 6*) (Fig. 4A–B). To test switch II involvement, we introduced alanine substitutions in Rap1b (Q63A, D69A, K73A) and assayed binding by immunoprecipitation. Rap1b^D69A^ emerged as a loss–of–function variant, supporting a switch II–mediated Rap1b–AC9 interaction (Fig. 4C–E). The phenotype was not due to misfolding: Rap1b^D69A^ still bound GTP and RalGDS–RBD, the latter mediated by switch I (Fig. 4D). Purified His–Rap1b^D69A^ failed to bind AC9–YFP by MST and to activate AC9 (Fig. 4E). Thus, AC9–C2a mediates interaction with Rap1b switch II. We superimposed the AC9 C1–C2:Rap1b model onto the AC9–Gαs cryo–EM structure, an overlay that guided identification of putative differential contact points for Gαs versus Rap1b (Fig. 4A–B). The model predicted Rap1b^D69^–AC9^Y1076^ contact specific to the Rap1b complex (Fig. 4F). We introduced alanine substitutions and quantified binding by MST (Fig. 4G). Likewise, AC9^Y1076A^–YFP lost binding to Rap1b but retained binding to Gαs (Fig. 4G). The overlay pinpointed an AC9^D1092^–Gαs^R280^ contact on the opposite face of the C2a cleft, a contact not observed in the Rap1b model (Fig.4F). MST confirmed that AC9^D1092A^ abolished Gαs binding while retaining Rap1b binding (Fig. 4G). Confocal studies confirmed that ISO drove AC9^WT^–YFP into a perinuclear compartment where it co–localized with mCherry–Rap1b^WT^ (Fig. S4); co–localization persisted with AC9^D1092A^ but was reduced with AC9^Y1076A^. Thus, AC9^Y1076A^ (Rap1b⁻, Gαs⁺) and AC9^D1092A^ (Rap1b⁺/Gαs⁻) discriminate AC9 binding partners.

**Fig. 4.**
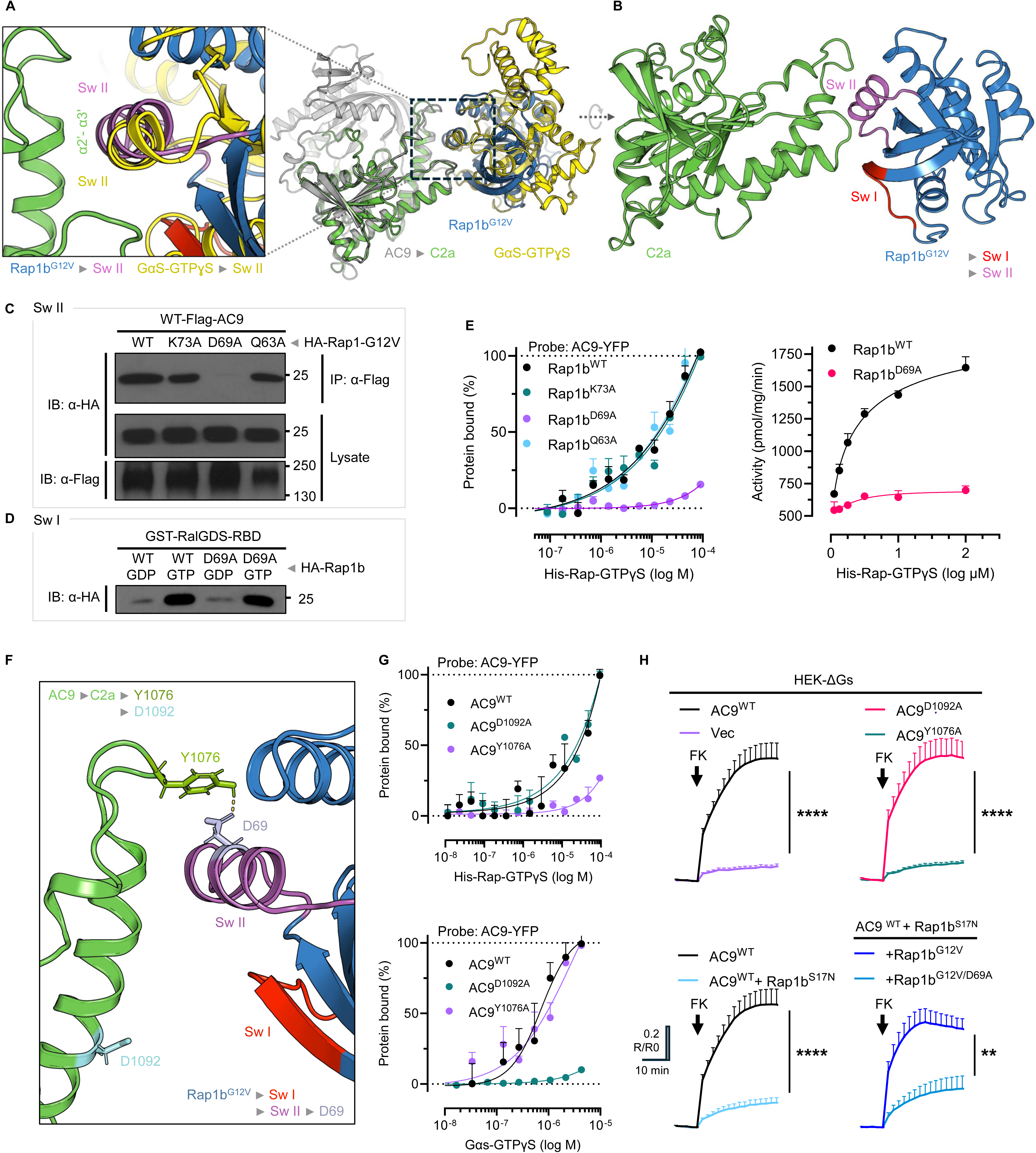
Rap1b switch II specifically engages the AC9 C2a cleft to enable isoform–specific activation. **(A, B)** AlphaFold–Multimer model of the AC9 C1–C2 catalytic domain bound to Rap1b^G12V^ (residues 1–166). (A) Overall view of the AC9 catalytic core (gray) with the C2a subdomain highlighted (green), showing Rap1b^G12V^ (blue) and Gαs–GTPγS (yellow) interacting with the C2a α2′–α3′ cleft. Close-up of the interface showing overlap of Rap1b switch II (purple) with the switch II region of Gαs–GTPγS (yellow) at the α2′–α3′ cleft of C2a (green) (left). (B) rotated/zoomed view (∼10°) of the Rap1b–C2a interface, highlighting Rap1b switch I (red) and switch II (purple). (**C**) Immunoprecipitation assays testing switch II involvement. Alanine substitutions within Rap1b switch II (Q63A, D69A, K73A) revealed Rap1b^D69A^ as a loss–of–function variant with reduced AC9 binding. (**D**) Rap1b^D69A^ preserved GTP loading and interaction with RalGDS–RBD (switch I–mediated), indicating that the reduced AC9 binding was not due to protein misfolding. (**E**) Microscale thermophoresis (MST) assay confirming loss of interaction between Rap1b^D69A^ and AC9 (left). *In vitro* AC9 activation assay showing that Rap1b^D69A^–GTP failed to activate AC9 (right). (**F**) Structural overlay of the AC9–Rap1b model with the AC9–Gαs cryo–EM structure, highlighting predicted differential contacts: Rap1b^D69^–AC9^Y1076^ specific to the Rap1b complex, and AC9^D1092^–Gαs^R280^ specific to the Gαs complex. (**G**) Differential AC9 mutant binding. AC9Y^1076A^–YFP (Rap1b⁻/Gαs⁺) lost binding to Rap1b but retained Gαs interaction, whereas AC9^D1092A^–YFP (Rap1b⁺/Gαs⁻) retained Rap1b binding but lost Gαs binding, as assessed by MST. (**H**) Real–time cAMP measurements using the H188 Förster resonance energy transfer (FRET) cAMP sensor to evaluate AC9 mutants functionally in HEK–ΔGαs cells. Traces represent normalized FRET ratios (R/R₀). Cells expressing the indicated AC9^WT^ or AC9 variants (AC9^D1092A^ or AC9^Y1076A^) were stimulated with forskolin (FK; 75 μM). Dominant-negative (Rap1b^N17^) or constitutively active Rap1b mutants (Rap1b^G12V^ or Rap1b^G12V/D69A^) were co-expressed with AC9^WT^ and H188. Traces are shown as mean ± SEM; n ≥ 8 cells from ≥ 3 independent experiments. Statistical significance was assessed at t = 20 min after FK addition (vertical line) using an unpaired two-tailed Student’s t test with Welch’s correction. Asterisks denote significance; **p < 0.0001, ****p < 0.0001. Structural models were rendered using PyMOL (panels A, B, and F).

We then turned to HEK–ΔGαs cells to evaluate AC9^WT^ alongside the AC9^Y1076A^ and AC9^D1092A^ variants (Fig.4H). AC9^D1092A^ (Rap1b⁺/Gαs⁻) reproduced the AC9^WT^ increase in FK responsiveness, whereas AC9^Y1076A^ (Rap1b⁻/Gαs⁺) attenuated this response. AC9^WT^ activation was inhibited by dominant–negative Rap1b^N17^ and rescued by Rap1b^G12V^, but not by Rap1b^G12V/D69A^, consistent with the switch II contact. Thus, in ΔGαs cells, the FK response depends on Rap1b–mediated activation of AC9, consistent with its CAP1–Rap1b sensitivity.

### Gαs–AC9 and Rap1b–AC9 control distinct phases of agonist–evoked cAMP signaling

Prompted by the compartment–specific effects of Rap1b–GTP on AC9 activity (Fig. 3D–G), we pondered whether this new AC regulation was involved in the endosomal ‘second’ cAMP wave. We first performed an orthogonal confirmation of the intracellular location of the AC9–Rap1b interaction. mAC9 also carries a split–TurboID(N) domain (*33*) (Fig. 5A–B), and upon co-expression of sTurboID(C)–Rap1b, TurboID reconstitution drives biotinylation that we monitored with the biotin sensor mSA–EYFP (34) as a proxy for the AC9–Rap1b interaction. As shown in Fig. 5A, in cells expressing mAC9 + sTurboID(C)–Rap1b^WT^, biotin alone or ISO alone rendered a diffuse mSA–EYFP signal; biotin + ISO triggered mSA–EYFP clustering exclusively at the intracellular/perinuclear AC9 pool, not at the PM. Endocytosis blockade abolished cluster formation, indicating that the Rap1b–accessible interface on AC9 emerges only after ISO activation and internalization (Figs. 5A–B).

**Fig. 5.**
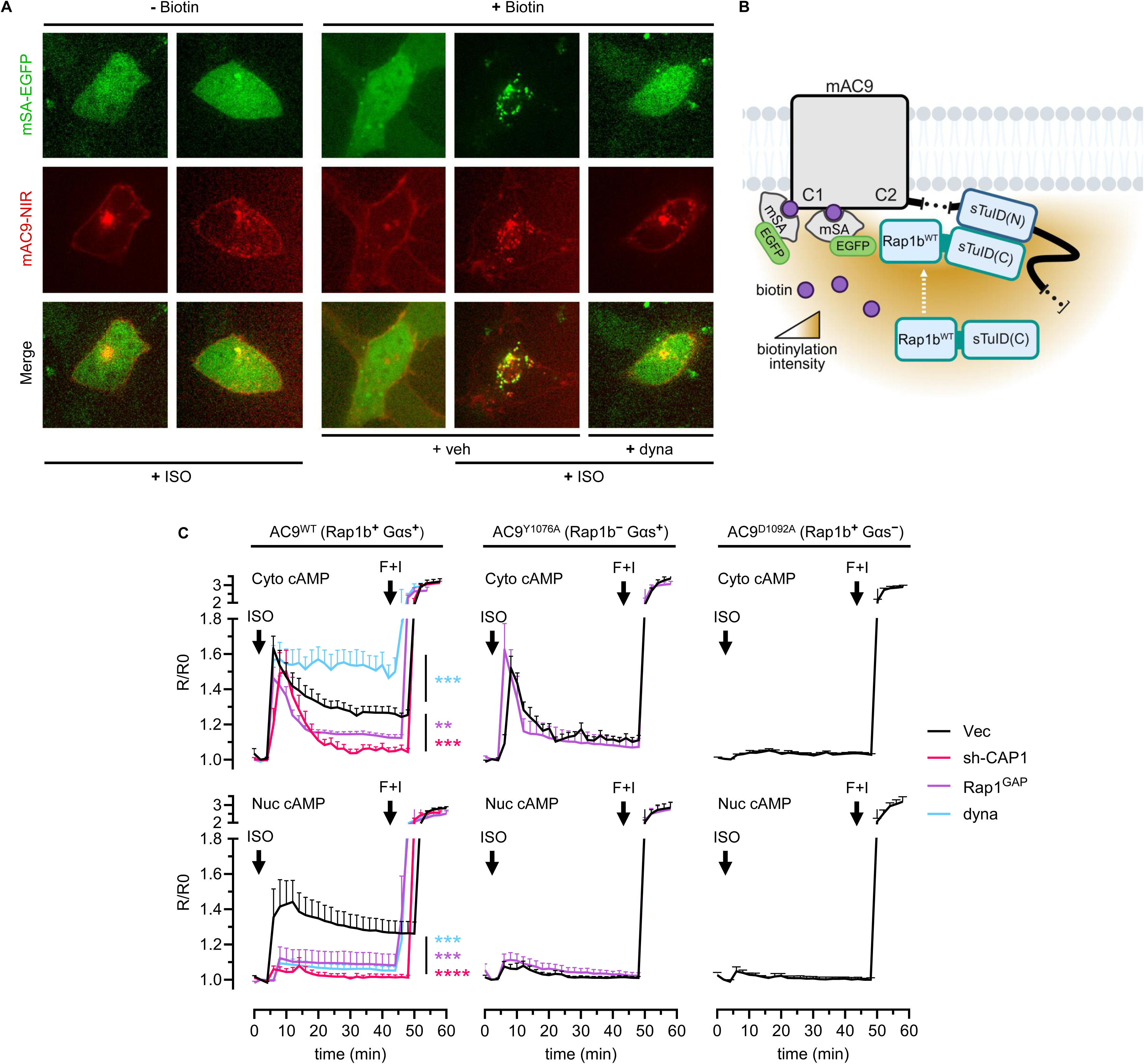
Compartment–specific regulation of AC9 by Gαs and Rap1b shapes sequential cAMP signaling waves. (**A**) Confocal micrographs of a split–TurboID proximity labeling assay in HC–1 cells coexpressing multi–AC9 (mAC9), Rap1bWT–sTurboID(C), and the biotin sensor mSA2–EYFP. Cells were treated with HBSS plus isoproterenol (−biotin + ISO) or with biotin (100μM) with or without isoproterenol (+ISO; 30nM), in the presence of either vehicle (veh; DMSO) or dynasore (dyna; 70μM) to block endocytosis. mAC9 contains the N–terminal half of split–TurboID [sTurboID(N)], whereas Rap1bWT is fused to the C–terminal half [sTurboID(C)]. Clustering of mSA2–EYFP indicates reconstituted TurboID activity and Rap1b–AC9 interaction. (**B**) Schematic of the split–TurboID proximity labeling assay. ISO–induced AC9–Rap1b interaction reconstitutes active TurboID at internalized AC9, leading to local biotinylation and mSA2–EYFP cluster formation. (**C**) Real–time cAMP measurements using the cytosolic FRET sensor H188 (top panels) or its nuclear–targeted version NLS–H188 (bottom panels) in live HC–1 cells co–expressing the sensors with AC9^WT^ (Rap1b⁺/Gαs⁺; left), AC9^Y1076A^ (Rap1b⁻/Gαs⁺; middle), or AC9^D1092A^ (Rap1b⁺/Gαs⁻; right). Cells were stimulated with ISO (30nM), followed by forskolin plus 3–isobutyl–1–methylxanthine (F+I; FK 20μM, IBMX 250μM). Traces are shown as mean ± SEM; n ≥ 7 cells from ≥ 3 independent experiments. Statistical significance was assessed at t = 20 min after ISO addition (vertical line) by one-way ANOVA with Dunnett’s multiple-comparisons test versus vector control. Color-matched asterisks denote significance (no asterisks, not significant); **p < 0.01, ***p < 0.001, ****p < 0.0001.

To assess the potential role of Rap1b–AC9 in the endosomal ‘second’ cAMP wave, we returned to HC–1 cells. AC9^WT^ triggered an ISO response that progressed from a PM first wave to a CAP1–Rap1b–dependent second wave after internalization (Fig. 5C), and both were required to generate nuclear cAMP (i.e., the ‘third wave’). Relative to WT, AC9^Y1076A^ (Rap1b⁻/Gαs⁺) left the first wave at the PM largely intact but markedly reduced the post–internalization sustained second wave. Consistent with this, the nuclear cAMP rise was absent. Removing CAP1 made AC9^WT^ behave like AC9^Y1076A^ (Rap1b⁻/Gαs⁺), underscoring the requirement for Rap1b. By contrast, AC9^D1092A^ (Gαs⁻/Rap1b⁺) eliminated both cytosolic and nuclear cAMP responses. Therefore, these combined results indicate that while Gαs–AC9 initiates and shapes the first cAMP wave, Rap1b–AC9 sustains a post–endocytic second wave. This Rap1b–CAP1–dependent second wave is indispensable for the subsequent nuclear cAMP wave and downstream transcription events.

## Discussion

In our previous work, we showed that the CAP1–Rap1b complex modulates cAMP dynamics in mammalian cells and defined how CAP1 independently engages Rap1b and ACs. We found that prenylated Rap1b inserts its geranylgeranyl group into the hydrophobic cavity of CAP1’s C–terminal β–solenoid, a nucleotide–independent interaction that leaves Rap1b’s switch regions available for other effectors (*19*). We also found that CAP1’s N–terminal coiled–coil domain binds all tested ACs through the C1a catalytic domain, leaving C2a available for binding of allosteric regulators (*20*). Together, these findings supported a scaffold model in which CAP1 positions Rap1b to bind and activate AC. In this study, we demonstrate that Rap1b, in complex with CAP1 but independent of Gαs, directly activates specifically the AC9 isoform.

The lack of AC isoform specificity of Rap1b–CAP1 *in vitro* (*20*) argues that AC9 selectivity in cells is driven primarily by additional regulatory constraints rather than by protein–domain affinity alone. We propose that the subcellular localization of AC9 is at least one key determinant of this specificity in cells. Because ligand–induced GPCR endocytosis is well characterized and Gαs–GTP is viewed as the canonical AC activator, most studies treat GPCR–G protein colocalization on intracellular membranes as a proxy for endomembrane signaling. This is typically assessed with conformation–selective nanobodies (*e.g*., Nb80, Nb37), Gαs–GTP–specific probes (*e.g*., KB1961), or engineered mini–Gs constructs (*35–43*). However, active GPCR–Gαs complexes neither guarantee colocalization with downstream ACs nor their activation, and the subcellular distribution and trafficking of individual AC isoforms are still largely unknown.

A limited number of studies have investigated AC endocytosis and found that AC9, but not AC1, undergoes internalization in response to ISO (*25, 44*). These findings are consistent with recent global organelle–profiling studies (*45*) that highlight AC9 as the only isoform with clear endosomal localization. Moreover, recent work suggests that GPCRs and Gαs can internalize through partially independent pathways, with GPCRs often reaching compartments that are deficient in G proteins (*46, 47*).

Although sustained Gαs–mediated signaling has been proposed for several GsPCRs, a PM–locked myristoylated Gαs still supports sustained cAMP production, raising the question of how AC9 is activated intracellularly (*48*). We propose that an endocytosis–dependent interaction between Rap1b and intracellular AC9, demonstrated by our biotinylation–proximity assay and by optogenetic recruitment of Rap1b^G12V^ studies, is sufficient to drive endosomal AC9 activation.

For Gαs, cryo–EM comparison of AC9 alone and in complex with Gαs–GTP shows that Gαs binding to the C2a domain remodels both C1a and C2a, including helix 4 near the FK site, to position active–site residues in an open, catalytically competent conformation. Although no Rap1b–AC9 structure is available, computational models predict that a helix from Rap1b switch II inserts into the C2a cleft between the α2′–α3′ helices, analogous to Gαs; our experimental validation of these models supports a conserved activation mechanism with allosteric communication to the FK site.

Here, we also identify and experimentally validate AC9 mutations that selectively disrupt Gαs–versus Rap1b–dependent binding and activation. In an AC–unresponsive background, introducing these mutants genetically uncouples Gαs- and Rap1b-mediated control of AC9, enabling dissection of their roles in ligand-evoked cAMP dynamics. These results support a sequence in which Gαs–AC generates the PM cAMP wave, followed by the Gαs-independent, Rap1b-dependent activation of endocytosed AC9 (second cAMP wave) that is necessary for the subsequent nuclear third wave within the three-wave framework (*18*).

Importantly, although Rap1b–AC9 is less efficacious than Gαs–AC9, Rap1b–GTP occupies a dual position in the pathway: it is generated downstream of cAMP as an Epac1 effector (*49, 50*), yet feeds back upstream by activating AC9 via CAP1, creating a positive feedback loop in which CAP1–Rap1b amplifies and sustains signaling. This provides a mechanism by which spatially restricted cAMP can relay information from transient PM Gαs–AC activity to sustained Rap1b–AC9 signaling in endocytic compartments. Where and how signaling switches from Gαs–AC to the Rap1b–AC9 complex, and how CAP1–Rap1b is targeted to AC9⁺ compartments, remain open questions.

In summary, we identify a Gαs-independent Rap1b–AC9 module that drives post-endocytic cAMP production by internalized AC9. To our knowledge, this work provides the first evidence that a small GTPase can activate a mammalian AC to generate the endosomal cAMP phase. Together with established PM Gαs–AC signaling and our prior identification of Ca²⁺-regulated nuclear sAC activity (9), these findings support a compartmentalized cAMP sequence that initiates at the cell surface, is relayed through endosomes, and culminates in the nucleus, with distinct regulatory control at each site. Extending our initial observations with TSHR (9) to the β₂AR, these results motivate future studies to determine how broadly this compartmentalized program applies across GPCRs.

## Materials and Methods

### Materials

Forskolin (F6886), IBMX (I7018), Dynasore (D7693), Isoproterenol (I6504), Digitonin (D141), Carbachol (212385), D–Biotin (B1595) were from Sigma–Aldrich (St. Louis, MO, USA). Furimazine (Nano–Glo Live Cell Assay System; N2011, Promega) was from Promega (Madison, WI, USA).

### Cell Lines and Transfections

The rat hepatoma clonal cell line HC–1 (kindly provided by Dr. Elliot Ross, University of Texas Southwestern Medical Center), human embryonic kidney WT HEK–293 cells (ATCC CRL–1573; Manassas, VA), and HEK–293 ΔGαs (kindly provided by Asaka Inoue, Tohoku University, Japan) cells were cultured in Dulbecco’s Modified Eagle’s Medium (DMEM; MT10013CM) supplemented with 10% fetal bovine serum (FBS; FB12999102), penicillin plus streptomycin (100 IU/mL and 100 μg/mL, respectively; 15140122), and L–glutamine (2 mM; 25030164) (All from Thermo Fisher Scientific). Cell lines were maintained at 37 °C in a humidified atmosphere containing 5% CO₂. Transient transfections were performed using Lipofectamine 3000 (L3000015; Thermo) for HC–1 cells or TurboFect Transfection Reagent (R0531; Thermo) for WT HEK–293 and HEK–293 ΔGαs cells. The total DNA amount was adjusted between 0.5 and 2 μg per well, according to the manufacturer’s instructions. To generate the HC–1 multi–AC9 stable cell line, cells were co–transfected with 2 μg of the multi–AC9 plasmid (see above) and 0.5 μg of pcDNA3.1–TA conferring G418 resistance. Forty–eight hours post–transfection, antibiotic selection was initiated and gradually increased to 800 μg/mL G418 (Geneticin; ant–gn–5; InvivoGen). over two weeks, with daily medium replacement. The surviving population was then sorted by flow cytometry (FACSAria III, BD Biosciences) to select the mAC9–expressing (NIR⁺) cells.

### Experimental solutions

Most experiments were performed in Hanks’ Balanced Salt Solution (HBSS; Gibco/Life Technologies, cat. no. 14185052) diluted to 1× in Milli–Q water and supplemented with CaCl₂ (1.8 mM), HEPES (20 mM), NaHCO₃ (4 mM), MgCl₂ (0.5 mM), and MgSO₄ (0.4 mM). The pH was adjusted to 7.4. All salts were from Sigma–Aldrich (St. Louis, MO, USA)

### Antibodies

Rabbit Monoclonal Antibody HA–Tag (C29F4) was from Cell Signaling (3724). His tag antibody (OAEA00010) was from Aviva Systems Biology. HA.11 antibody (901513) was from Biolegend, Inc. Mouse monoclonal antibody anti–Flag M2 (F1804), Donkey Anti–Rabbit Alexa–488 (A21206), Goat Anti–Mouse Alexa–555 (A32727) were from Sigma–Aldrich (St. Louis, MO, USA)

### DNA Constructs

For a complete list of constructs used in this study, see Table S1. AC2/AC9 chimeric adenylyl cyclases. A panel of six AC2/AC9 chimeric adenylyl cyclases was designed in–house and synthesized by GenScript (USA). Each construct contained an N–terminal HA epitope tag and consisted of domain–swapped fragments between rat AC2–HA and human Flag–AC9 (Table S1). Multi–tagged AC9 (mAC9). mAC9 was derived from bovine AC9–YFP by removing the YFP coding sequence and inserting an in–frame multifunctional cassette (see Fig. S3 for a schematic representation). The cassette included: eMagA^F^ (enhanced ‘Magnets’ photodimerization module for blue–light–dependent heterodimerization; used with the complementary eMagB^F^ module; sTurboID(N) (low–affinity N–terminal split–TurboID fragment, residues 2–73; Q65P; split junction at L73; Addgene #153002) for complementation–dependent proximity biotinylation with sTurboID(C); mNG2(11) (16–aa peptide corresponding to residues 214–229 of mNeonGreen2; split between the 10th and 11th β–strands and removal of the additional GFP–like C terminus GMDELYK, yielding a 213–aa mNG1–10 and a 16–aa mNG11) to be used with C–terminally fused constructs for complementation with mNG2(1–10); HiBiT or SmBiT for luciferase complementation with LgBiT; miRFP670nano3 (GenBank MW627294.1) for near–infrared optical detection; and HA and ALFA epitope tags for antibody–/nanobody–based detection and affinity capture. The following sequences were used: ALFA tag (DNA): 5′–TCACGTTTGGAAGAGGAACTGAGACGCCGCTTAACTGAA–3′; SmBiT (DNA): 5′–GTGACCGGCTACCGCCTGTTCGAGGAGATCCTG–3′; HA tag (peptide): YPYDVPDYA.

Flexible linkers were inserted between modules (e.g., GGS, GGSG, SGGGGS, GNSGSSGGGGSGGGGSSG, [G4S]n, KGSGSTSGSG).

### Bacterial expression constructs

His–C1a (aa 380–574) and His–C2a (aa 1049–1245) from AC9 were generated by PCR using AC9–YFP as template. PCR products were digested with NdeI/XhoI and subcloned into pET–28c. Primers were:

- C1a FW: 5′–GTCAGTCACATATGATCGCCTTCCGGCCCTTCAAGATGCAGCAGAT–3′
- C1a REV: 5′–GTCAGTCACTCGAGTTACTGGCCGGCGATCAGGTAGGTTTTCAG–3′
- C2a FW: 5′–AGCTAGCTCATATGCAGACCTACAGCAAGAACCACGAC–3′
- C2a REV: 5′–AGCTAGCTCTCGAGTTACTTGGGGTACAGGTATGTCTTCAT–3′

His–GFP–RapG12V was generated by subcloning GFP–Rap1 into pET–28c using NcoI/BamHI sites. Primers were:

- FW: 5′–AGCTCCATGGGCAGCAGCCATCATCATCATCATCACGTGAGCAAGGGCGAGGAGCTG–3′
- REV: 5′–AGCTGGATCCTTAAAGCAGCTGACATGATGACTT–3′

Unless otherwise noted, additional modified AC constructs, engineered derivatives, and other constructs were generated by de *novo* DNA synthesis (GenScript) followed by assembly into pcDNA3.1 under the cytomegalovirus (CMV) enhancer–promoter. The positions and order of inserted modules, amino–acid substitutions, and other modifications are summarized in Table S1. All plasmids were confirmed by Sanger sequencing (full ORF and all cloning junctions).

### Microscopy

Widefield. Imaging was performed on an Olympus IX83 motorized two–deck inverted microscope equipped with a six–line multi–LED Spectra X light engine (Lumencor), a Prior emission filter wheel, and a Prior ProScan XY stage (Prior Scientific), using a Teledyne Photometrics Kinetix back–illuminated sCMOS camera (01–KINETIX–M–C; 10.2 MP; 3200 × 3200 pixels; 6.5–µm pixels; 20.8 × 20.8–mm sensor; rolling shutter; 16–bit). The microscope was controlled with SlideBook 6 (Intelligent Imaging Innovations). Confocal. The system was fitted with an X–Light V3 spinning disk confocal unit (CrestOptics; lateral/axial resolution [FWHM] ∼230 nm/∼600 nm) equipped with an LDI–7 multiline solid–state laser illuminator (89 North). For FRET measurements (widefield), images were acquired using 4×4 binning in dynamic range mode (conversion gain, 0.23) with either a LUCPlanFL N 20×/0.45 NA air objective (∞/0–2/FN26.5) or a UPlanXApo 40×/1.4 NA oil objective (∞/0.17/FN26.5). For optogenetic stimulation (see below), immunofluorescence, and other confocal experiments, images were acquired using 2×2 binning in sub–electron mode (conversion gain, 0.015) with a UPlanSApo 100×/1.4 NA oil objective (∞/0.17/FN26.5).

### FRET–based cAMP measurements

Cells expressing the H188 or NLS–H188 sensor were seeded onto 18–well chambered slides (ibidi µ–Slide 18–Well Chambered Coverslip, 81816) or 96–well round µ–plates with #1.5H high–precision glass bottoms (ibidi, 89626). After several washes with HBSS, cells at ∼80% confluence (≥48 h post–transfection) were imaged on the setup described above (widefield mode). The H188 sensor was excited at 440 nm (20% intensity), and fluorescence emission was collected through 470/30 nm (donor) and 535/30 nm (acceptor) filters using an 86002v1 dichroic (Chroma Technology Corp.). Signals were background–subtracted and corrected for donor bleed–through into the acceptor channel (∼53%) and direct acceptor excitation (∼3%). No significant photobleaching was detected under these acquisition conditions. Isoproterenol (ISO) stimulation was analyzed by calculating FRET ratios (R; 470/535) normalized to the pre–stimulus baseline (R/R₀). Unless otherwise indicated, responses were quantified 20 min after ISO addition. At the end of each experiment, forskolin (FSK; 20 µM) plus IBMX (250 µM) were added to elicit the maximal sensor response and verify that the sensor was not saturated during ISO stimulation. For ISO dose–response experiments (DR protocol), each ISO dose was tested in separate wells/cells to capture both peak and steady–state responses. FRET ratios (R/R₀) were normalized to baseline (no agonist; 0%) and to the response elicited by a saturating ISO concentration (1000 nM, 20 min; 100%).

### Optogenetic stimulation and local cAMP imaging

Optogenetic stimulation was delivered with 445–nm light from a solid–state laser illuminator (LDI–7, 89 North) coupled to a point–scanning module (UGA–42 Geo, Rapp OptoElectronic) mounted on a backport of the confocal setup, enabling patterned/focal photostimulation while simultaneously acquiring FRET/intensiometric signals through an independent imaging light path. Stimulation was applied at 10% laser power with ND 10% neutral–density attenuation, routed through a 458–nm longpass dichroic (ZT458rdc, Chroma). Circular stimulation ROIs (radius 5 scanner units; ∼1 µm²) were illuminated at predefined subcellular locations for 50 cycles at 4 Hz (250–ms period; 200 ms ON/50 ms OFF; total train 12.5 s; cumulative illumination 10.0 s; 80% duty cycle). HC–1 cells stably expressing multi–tagged AC9 (mAC9; containing eMagA^F^ and an ALFA tag) were transiently transfected with eMagB^F^–Rap1b^G12V^ and the NbALFA–H208 FRET cAMP sensor, which is tethered to mAC9 via ALFA binding to report local cAMP near AC9. To minimize unintended photostimulation and light–associated phototoxicity, cells expressing optogenetic constructs were handled under low–blue safelight illumination (GBX–2 dark red filter, 5.5″; 13–W amber compact fluorescent bulb; Low Blue Lights). Cells were imaged 48 h post–transfection. For each cell, seven spatially contiguous stimulation ROIs were illuminated to target mAC9 at the plasma membrane, intracellular mAC9–positive clusters, cytosolic regions lacking mAC9, and an extracellular region. Local cAMP responses were quantified from three predefined acquisition ROIs: a plasma–membrane segment, a perinuclear region, and the whole–cell (global) area (Fig. 3).

### Immunofluorescence staining

Cells were grown on glass coverslips in 6–well plates. For staining, cells were rinsed with phosphate–buffered saline (PBS, pH 7.4), fixed in 4% paraformaldehyde for 15 min at room temperature, and permeabilized with 0.5% Triton X–100 in PBS for 15 min. Cells were blocked in PBS containing 3% bovine serum albumin (BSA), followed by incubation with primary antibodies. Coverslips were washed 3–5 times (5 min each) with PBS and incubated with secondary antibodies. After 3–5 additional washes (5 min each), coverslips were mounted in ProLong Glass Antifade Mountant (P36984, Thermo) and imaged using a setup described above (confocal mode) with appropriate filter settings. Images were analyzed using SlideBook 6 software, ImageJ, and Icy software.

### Monitoring of Split–TurboID–Mediated Biotinylation

To monitor sTurboID–catalyzed biotinylation in living cells, we used an engineered monomeric streptavidin (K56R) fused to enhanced green fluorescent protein (mSA–EYFP) as a fluorescent reporter (33, 34). Stable HC–1 mAC9 cells were transiently transfected with mSA–EYFP and sTurboID(C)–bRap1b^WT^, which interacts with the sTurboID(N) domain of mAC9. After transfection, cells were maintained under nominally biotin–free conditions in high–glucose DMEM (Gibco 11965, Thermo Fisher Scientific) supplemented with charcoal–stripped FBS (A3382101, Thermo Fisher Scientific). To further reduce residual low–molecular–weight biotin, FBS was diafiltered using 10 kDa Amicon Ultra–15 centrifugal filters by repeated concentrate–and–reconstitute cycles (serum concentrated by centrifugation, filtrate discarded, and the retentate restored to the starting volume with sterile 1× PBS). Before imaging, cells were washed in HBSS and baseline fluorescence was recorded; D–biotin was then added to a final concentration of 100 µM to initiate labeling and monitor biotinylation–dependent mSA–EYFP redistribution, in the absence or presence of isoproterenol (30 nM).

### Protein purification

For *E. coli* expression, Rosetta (DE3) cells (Novagen, 71403–3) were transformed with the indicated pGEX or pET28c plasmids were grown to an OD₆₀₀ > 1.0 and induced with 0.5 mM IPTG for 16 h at 24 °C. Cells were harvested and lysed in buffer containing 50 mM Tris (pH 7.5), 150 mM NaCl, 10% glycerol, 1% Triton X–100 (Sigma–Aldrich, X114), 0.2 mg/mL lysozyme, and PMSF. Lysates were clarified by centrifugation, and supernatants were subjected to affinity purification on GSH–Sepharose (4B GST–tagged protein purification resin; Cytiva, 17075605) or Ni–NTA agarose (Qiagen, 30210), as appropriate.

Gαs–8×His expressed in High Five insect cells was solubilized in 1% dodecyl–β–maltoside (DDM, Cayman, 16494) and purified using Ni–NTA agarose. Nucleotide loading was performed by incubating purified proteins with 0.1 mM GTPγS and 10 mM MgCl₂ at 30 °C for 1h. Mammalian AC9 was purified by affinity chromatography from stably AC9-3C-YFP-twinStrep-transfected HEK cells, as previously described (6). Briefly, after a hypotonic lysis step, a membrane fraction was prepared by ultracentrifugation and solubilized in a buffer containing 1% DDM and 0.2% cholesteryl hemisuccinate (CHS) at 4°C for 12h. A clarified lysate was then mixed with Strep-Tactin beads for 1h, washed extensively in a buffer containing 0.1% digitonin, and proteins eluted by incubating with HRV3C protease (AC9), or 500 mM biotin (AC9-YFP), overnight at 4°C. Upon concentration (Amicon Ultra centrifugal filter unit; 50 kDa MWCO, MilliporeSigma), samples were further purified by size-exclusion chromatography (Superose 6 Increase 10/300 GL column; Cytiva).

### *In vitro* AC9 Activation

Purified AC9 or AC9-YFP (0.1 μg) was mixed with 100 nM Gαs-GTPγS in assay buffer (100µl; 50 mM Tris, pH 8.0, 5 mM MgCl₂, 1 mM ATP) and incubated for 30 min at 30°C. Reactions were stopped (100°C for 10 min), followed by centrifugation at 15,000 rpm for 10 min. Supernatants were collected, and aliquots were assayed using the Cyclic AMP XP Assay Kit (4339, Cell Signaling Technology), according to the manufacturer’s instructions. For Rap1 displacement, purified His-Rap1-GTPγS (0 to 6µM) was included in the assay. Forskolin dose-responses were performed under the same assay conditions utilizing purified AC9 (0.1 µg) and His-Rap1-GTPγS or His-Rap1-GDP (2 µM).

### Immunoprecipitation

HEK–293 cells were co–transfected with Flag–AC9 and either HA–Rap1b^G12V^, HA–Rap1b^K73A^, HA–Rap1b^D69A^, or HA–Rap1b^Q63A^. Forty–eight hours after transfection, cells were lysed in buffer containing 50 mM Tris (pH 7.5), 50 mM NaCl, 0.5% Nonidet P–40, 5% glycerol, 5 mM MgCl₂, 1 mM GTPγS, and protease inhibitors (539131–10VL, Sigma). Lysates were incubated for 1 h at 4 °C with Flag–agarose beads (M8823, Millipore), followed by four washes in lysis buffer. Bound proteins were eluted in 2% SDS and analyzed by immunoblotting with anti–Flag or anti–HA antibodies.

### Microscale thermophoresis (MST) assays

Measurements were performed on a Monolith NT.115 instrument (NanoTemper) using Premium–coated capillaries (NanoTemper, MO–K025). Experiments were conducted using 20–80% LED power and medium or high MST power. The MST binding buffer contained 50 mM Tris–HCl (pH 8.0), 150 mM NaCl, 5 mM MgCl₂, 0.1 mM GTPγS (or GDP), and 0.02% digitonin. Dose–response curves were acquired in triplicate (mean ± SEM) and analyzed using the instrument software or after import into GraphPad Prism 10.

### Rap1 Activation Assay Using RalGDS–RBD

Cells expressing HA–Rap1b or HA–Rap1b–D69A were lysed in buffer containing 25 mM Tris–HCl, pH 7.5, 150 mM NaCl, 1% Nonidet P–40, 5% glycerol, 5 mM MgCl₂, and protease inhibitors. Lysates were clarified by centrifugation at 13,000 rpm for 10 min at 4 °C. For each 500 μL lysate, 10 μL of 0.5 M EDTA (final 10 mM) was added, and samples were vortexed. GTPγS (5 μL of 10 mM; final 0.1 mM) or GDP (5 μL of 100 mM; final 1 mM) was then added, vortexed, and the mixtures were incubated at 30 °C for 30 min with gentle agitation. Reactions were stopped by placing samples on ice and adding 32 μL of 1 M MgCl₂ (final 60 mM). Purified GST–RalGDS–RBD bound to glutathione–Sepharose beads (10 μg) was added and incubated with the supernatants for 60 min at 4 °C with rotation. Beads were washed four times with lysis buffer, resuspended in Laemmli sample buffer (1610737, Bio–Rad), and analyzed by SDS–PAGE (12%) followed by transfer to PVDF membranes (EMD Millipore, IPVH00010) for immunoblotting.

### His Pull–Downs

His–tagged proteins (His–GFP, His–C1a, and His–C2a; 50 μg) were expressed in *E. coli* and captured on 40 μL of 50% Ni–NTA agarose beads (Qiagen, 30210) by rotation at 4 °C for 1 h. Beads were briefly centrifuged and washed three times with lysis buffer (25 mM Tris–HCl, pH 7.5, 150 mM NaCl, 5 mM MgCl₂, 5% glycerol, 1% Nonidet P–40). Lysates from HEK293 cells transfected with HA–Rap1^G12V^ were prepared in the same buffer supplemented with 0.1 mM GTPγS. Lysate (5 μL; 5–10 μg/μL) was added to the beads in 800 μL binding buffer and incubated for 1 h at 4 °C with rotation. Beads were washed three times, and bound proteins were eluted, resolved by SDS–PAGE, and analyzed by immunoblotting with an anti–HA antibody.

### Statistics

Dose–response curves were fit by nonlinear regression (four–parameter logistic, variable slope) in GraphPad Prism 10. EC₅₀ values (and Hill slopes) were obtained from fits to each independent experiment and are reported with 95% confidence intervals. For comparisons of dose–response curves, extra–sum–of–squares F tests were used where indicated. For comparisons between two groups, two–tailed unpaired Student’s t tests (Welch’s correction when appropriate) were used; for ≥3 groups, one–way ANOVA with Dunnett’s multiple–comparisons test versus vector control was used (α = 0.05). Data are presented as mean ± SEM unless otherwise indicated. n denotes independent experiments unless stated otherwise.

## Acknowledgments

We thank Drs. Elliott M. Ross (University of Texas Southwestern Medical Center) and Asuka Inoue (Tohoku University, Japan) for providing HC-1 and Gαs-knockout cells, respectively. We thank Drs. Volodymyr Korkhov (Paul Scherrer Institute, Switzerland), Dermot Cooper (University of Cambridge), and Carmen W. Dessauer (UTHealth Houston, McGovern Medical School) for sharing adenylyl cyclase constructs and related reagents. We thank Dr. Philip J. S. Stork (Oregon Health & Science University) for mCherry–Rap1 constructs and Dr. Yubin Zhou (Texas A&M University) for the mSA2–EYFP construct. We thank Dr. Lutz Birnbaumer (Instituto de Investigaciones Biomédicas, UCA, Argentina) for critical reading of the manuscript.

## Author contributions

Conceptualization: DLA, AP^1^, XZ AP^2^

Methodology: XZ, AP^1^, AP^2^

Funding acquisition: DLA

Project administration: DLA

Supervision: DLA

Writing – original draft: DLA, AP^1^

Writing – review & editing: DLA, AP^1^, AP^2^

^1^Alejandro Pizzoni;

^2^Alipio Pinto

## Data and materials availability

All data are available in the main text or the suppl. materials.

## Supplementary Materials

Figs. S1 to S4

Table S1

