## Supplementary material for "Rap1b Activates Endosomal AC9 to Drive the Second cAMP Wave": Figs. S1 to S4; Table S1

**The PDF file includes:**

Figs. S1 to S4
Table S1

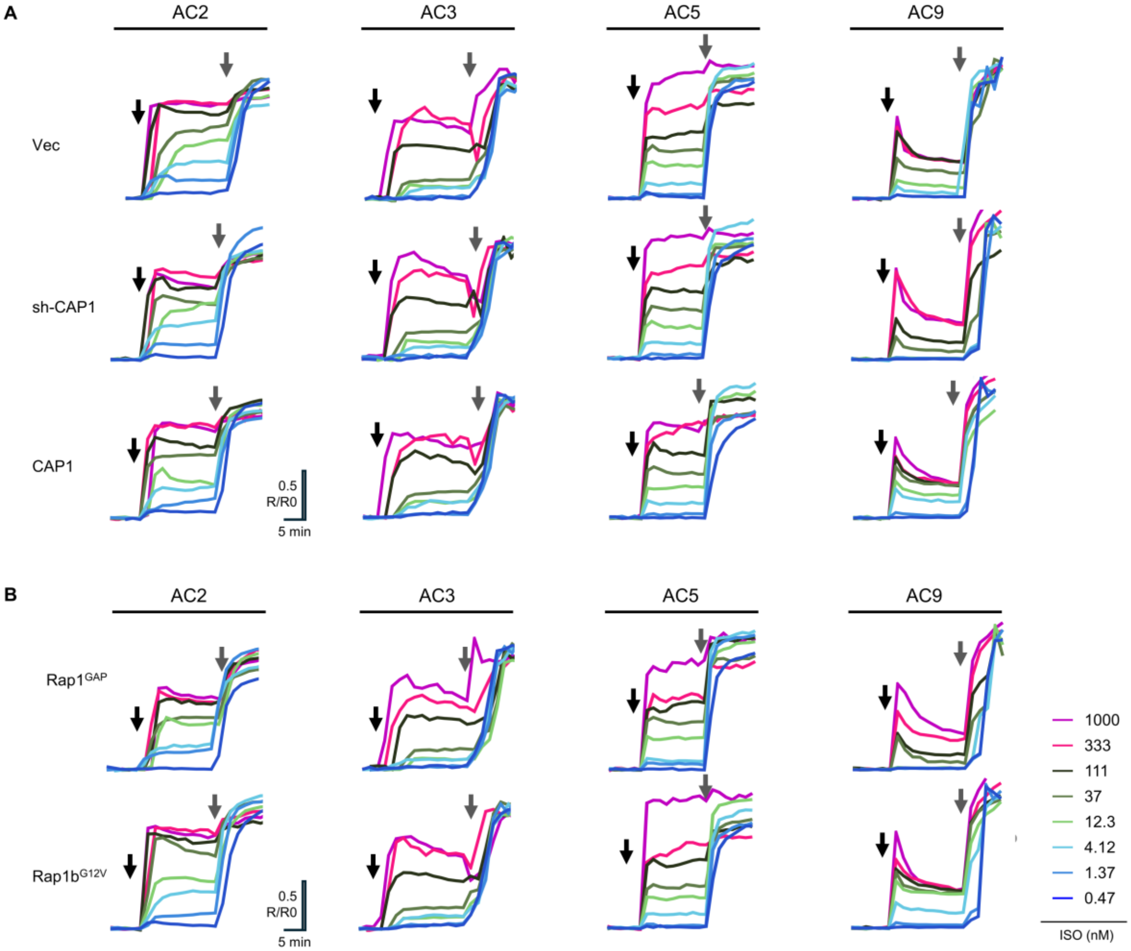


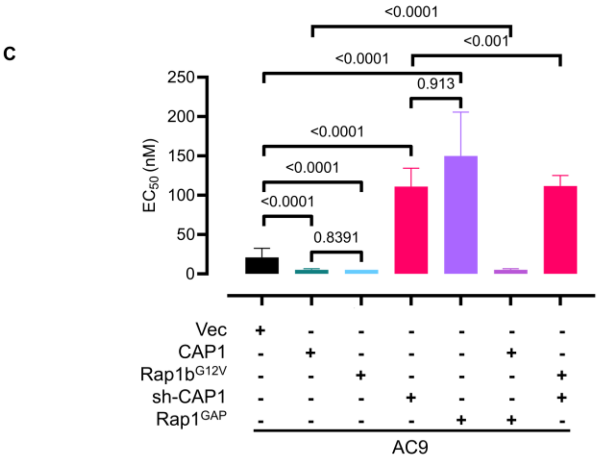


| | $EC_{50}$ | $EC_{50}$ 95% CI lo | $EC_{50}$ 95% CI hi |
|---|---|---|---|
| **Vec** | 20.86 | 13.2 | 32.46 |
| **sh-CAP1** | 111 | 90.43 | 134.4 |
| **CAP1** | 5.131 | 4.061 | 6.55 |
| **Rap1GAP** | 149.8 | 107.8 | 205.7 |
| **Rap1bG12V** | 5.106 | 3.476 | 7.564 |
| **CAP1+Rap1GAP** | 5.131 | 4.061 | 6.55 |
| **Rap1bG12+sh-CAP1** | 111.6 | 99.59 | 125 |

**Fig. S1 (**A–C**).** Isoform–specific cAMP responses to isoproterenol (ISO) stimulation in HC–1 cells. Real–time FRET measurements of cytosolic cAMP using the H188 sensor in live HC–1 cells expressing different AC isoforms (AC2, AC3, AC5, or AC9). Traces show mean normalized FRET ratios ($R/R_0$) in response to increasing concentrations of isoproterenol (ISO; 0.47–1000 nM; black arrow); SEM is omitted for visual clarity. After ISO, cells were stimulated with forskolin plus IBMX (FK, 20 μM; IBMX, 250 μM; gray arrow). For AC3–expressing cells, recordings were performed in the presence of carbachol (150 μM), as required for robust AC3 activation. (**A**) Effect of CAP1 overexpression (CAP1) or knockdown (sh–CAP1) on cAMP production. (**B**) Effect of Rap1b modulation using constitutively active Rap1bG12V or Rap1GAP–mediated Rap1b inactivation on cAMP production. Traces in panels A and B were used to generate the dose–response (DR) curves in Fig. 1A and 1B. (**C**) $EC_{50}$ values were obtained by nonlinear regression fits of the dose–response curves in Fig. 1A, 1B, and 1C to a four–parameter logistic equation, with 95% confidence intervals (CI). Differences between curves were assessed by extra–sum–of–squares F tests, and the corresponding p values are shown above the pairwise comparisons. (**D**) All data are representative of at least 3 independent experiments. *Figure S1 continues on the following page.*

D

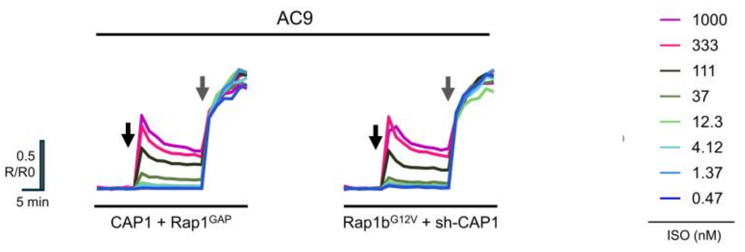
AC9
0.5
R/R0
5 min
CAP1 + Rap1GAP
Rap1bG12V + sh-CAP1
1000
333
111
37
12.3
4.12
1.37
0.47
ISO (nM)

E

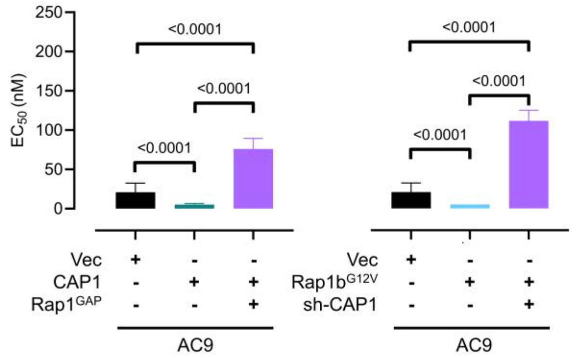
EC50 (nM)
250
200
150
100
50
0
<0.0001
<0.0001
<0.0001
<0.0001
<0.0001
<0.0001
Vec + - -
CAP1 - + +
Rap1GAP - - +
AC9
Vec + - -
Rap1bG12V - + +
sh-CAP1 - - +
AC9

F

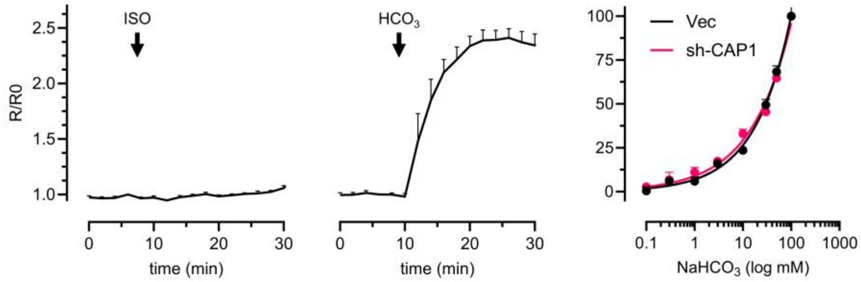
R/R0
2.5
2.0
1.5
1.0
ISO
HCO3
time (min)
0 10 20 30
time (min)
0 10 20 30
Vec
sh-CAP1
100
75
50
25
0
0.1 1 10 100 1000
NaHCO3 (log mM)

G

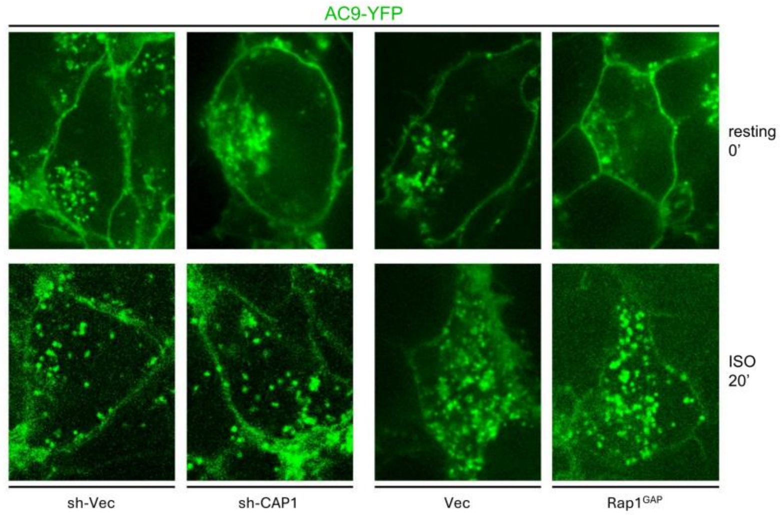
AC9-YFP
resting
0'
ISO
20'
sh-Vec
sh-CAP1
Vec
Rap1GAP

**Fig. S1, continued (D–G).**

(**D**) **Combined CAP1–Rap1b modulation in HC–1 cells expressing AC9.** Real–time FRET measurements of cytosolic cAMP using the H188 sensor in live HC–1 cells expressing AC9. Traces show mean normalized FRET ratios ($R/R_0$) in response to increasing concentrations of isoproterenol (ISO; 0.47–1000 nM; black arrow); SEM is omitted for visual clarity. After ISO, cells were stimulated with forskolin plus IBMX (FK, 20 µM; IBMX, 250 µM; gray arrow). Left: CAP1 overexpression together with $Rap1^{GAP}$. Right: sh–CAP1 together with constitutively active $Rap1b^{G12V}$. Traces in panel D were used to generate the dose–response (DR) curves in Fig. 1C. (**E**) $EC_{50}$ values were obtained by nonlinear regression fits of the dose–response curves in Fig. 1C to a four–parameter logistic equation, with 95% confidence intervals (CI). Differences between curves were assessed by extra–sum–of–squares F tests, and the corresponding p-values are shown above the pairwise comparisons. (**F**) **HC–1 cells are insensitive to ISO and exhibit an endogenous CAP1–insensitive sAC activity.** Real–time FRET measurements of cytosolic cAMP (H188 sensor) in HC–1 cells reveal no detectable response to isoproterenol (ISO; 100 nM; left panel), while showing robust activation of soluble AC (sAC), evidenced by a marked cAMP increase upon $HCO_3^-$ stimulation (100 mM; middle panel). Dose–response curves of $HCO_3^-$–evoked cAMP production in HC–1 cells transfected with vector control or sh–CAP1. (**G) CAP1 and Rap1b manipulations do not alter ISO–induced AC9 internalization.** Confocal micrographs of HC–1 cells expressing AC9–YFP under four conditions: sh–Vector, sh–CAP1, Vector, or $Rap1b^{GAP}$. The top row shows the basal distribution of AC9–YFP, and the bottom row shows AC9–YFP after 20 min of stimulation with isoproterenol (ISO; 30 nM). Cells were fixed either at rest or after ISO stimulation. In all conditions, the resting state shows the intracellular perinuclear cluster normally observed for this isoform. Upon ISO stimulation, AC9 undergoes robust agonist–induced internalization, indicating that neither CAP1 depletion nor Rap1b inactivation detectably alters AC9 trafficking. All data are representative of at least 3 independent experiments.

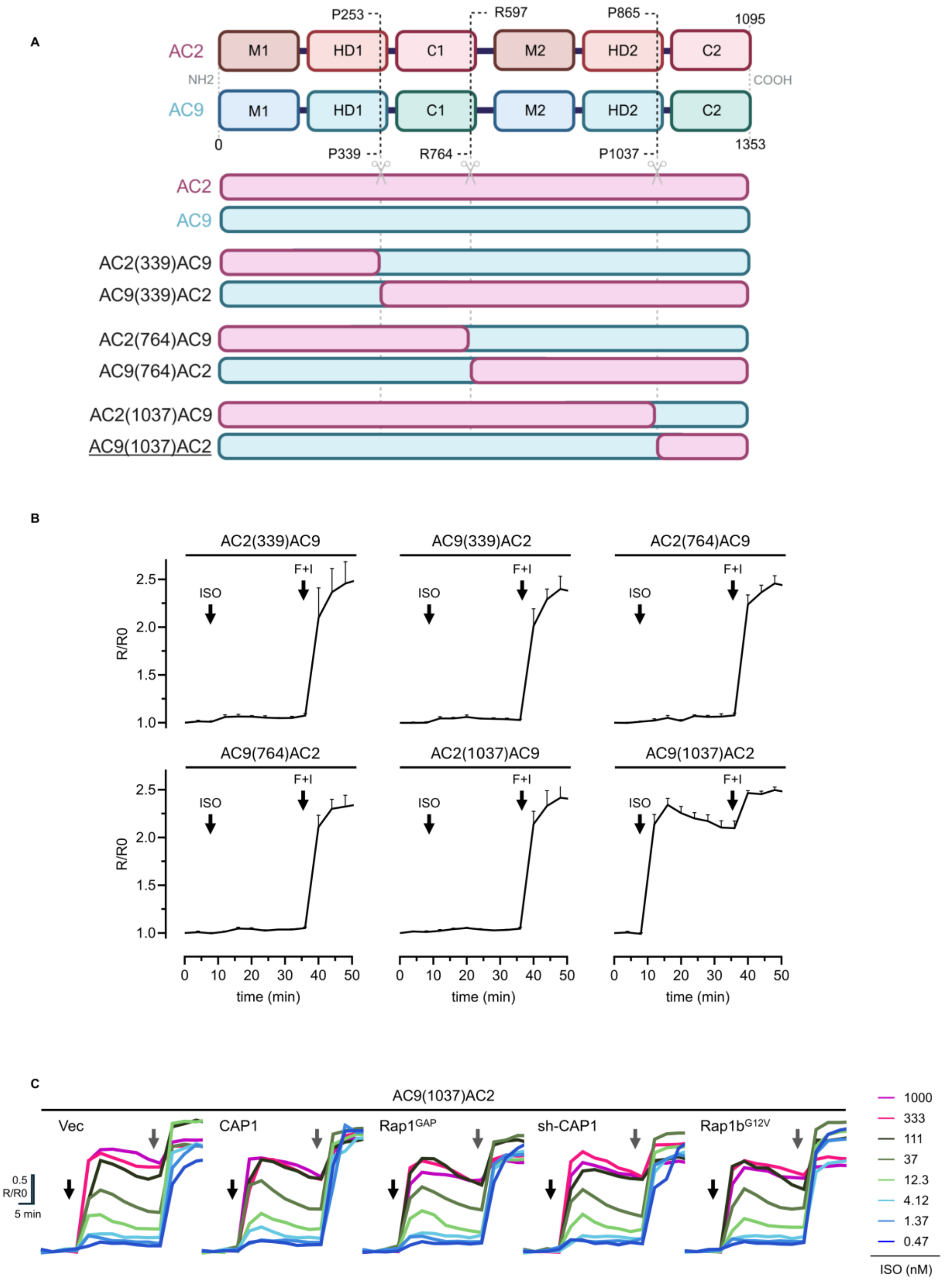
A
AC2
AC9
M1
HD1
C1
M2
HD2
C2
P253
R597
P865
1095
NH2
COOH
0
1353
P339
R764
P1037
AC2(339)AC9
AC9(339)AC2
AC2(764)AC9
AC9(764)AC2
AC2(1037)AC9
AC9(1037)AC2
B
ISO
F+I
R/R0
time (min)
AC9(1037)AC2
C
Vec
CAP1
Rap1GAP
sh-CAP1
Rap1bG12V
0.5 R/R0
5 min
1000
333
111
37
12.3
4.12
1.37
0.47
ISO (nM)

**Fig. S2** (A–C)**. Domain organization, design, and functional validation of AC2/AC9 chimeric constructs.** (**A**) Schematic representations of the domain architecture of AC2 and AC9 illustrating the regions selected to generate six AC2/AC9 chimeric constructs, each combining defined segments from the two ACs. Amino acid positions corresponding to the junctions between AC2– and AC9–derived regions are indicated by dotted lines for all chimeras. Each chimera is named according to the AC9 junction position, whereas the corresponding AC2 junction is depicted in the cartoon. M1/M2, transmembrane domains 1 and 2; HD1/HD2, helical domains 1 and 2; C1/C2, catalytic domains 1 and 2. (**B**) Functional characterization of AC2/AC9 chimeras identifies AC9(1037)AC2 as the only active construct. Real–time FRET measurements of cytosolic cAMP using the H188 sensor in HC–1 cells expressing each of the six AC2/AC9 chimeric ACs. Traces represent mean normalized FRET ratios ($R/R_0$) in response to isoproterenol (ISO; 1000 nM) followed by FK plus IBMX (F+I; FK 20 μM, IBMX 250 μM). (**C**) Evaluation of AC9(1037)AC2 sensitivity to CAP1 and Rap1b manipulations. Real–time FRET measurements of cytosolic cAMP using the H188 sensor in HC–1 cells expressing AC9(1037)AC2 and either Vector, CAP1, $Rap1^{GAP}$, sh–CAP1, or $Rap1b^{G12V}$. Traces show mean normalized FRET ratios ($R/R_0$) in response to increasing concentrations of isoproterenol (ISO; 0.47–1000 nM; black arrow); SEM is omitted for visual clarity. After ISO, cells were stimulated with forskolin plus IBMX (FK, 20 μM; IBMX, 250 μM; gray arrow). Traces were used to generate the dose–response (DR) curves in Fig. 2D. All data are representative of at least 3 independent experiments.

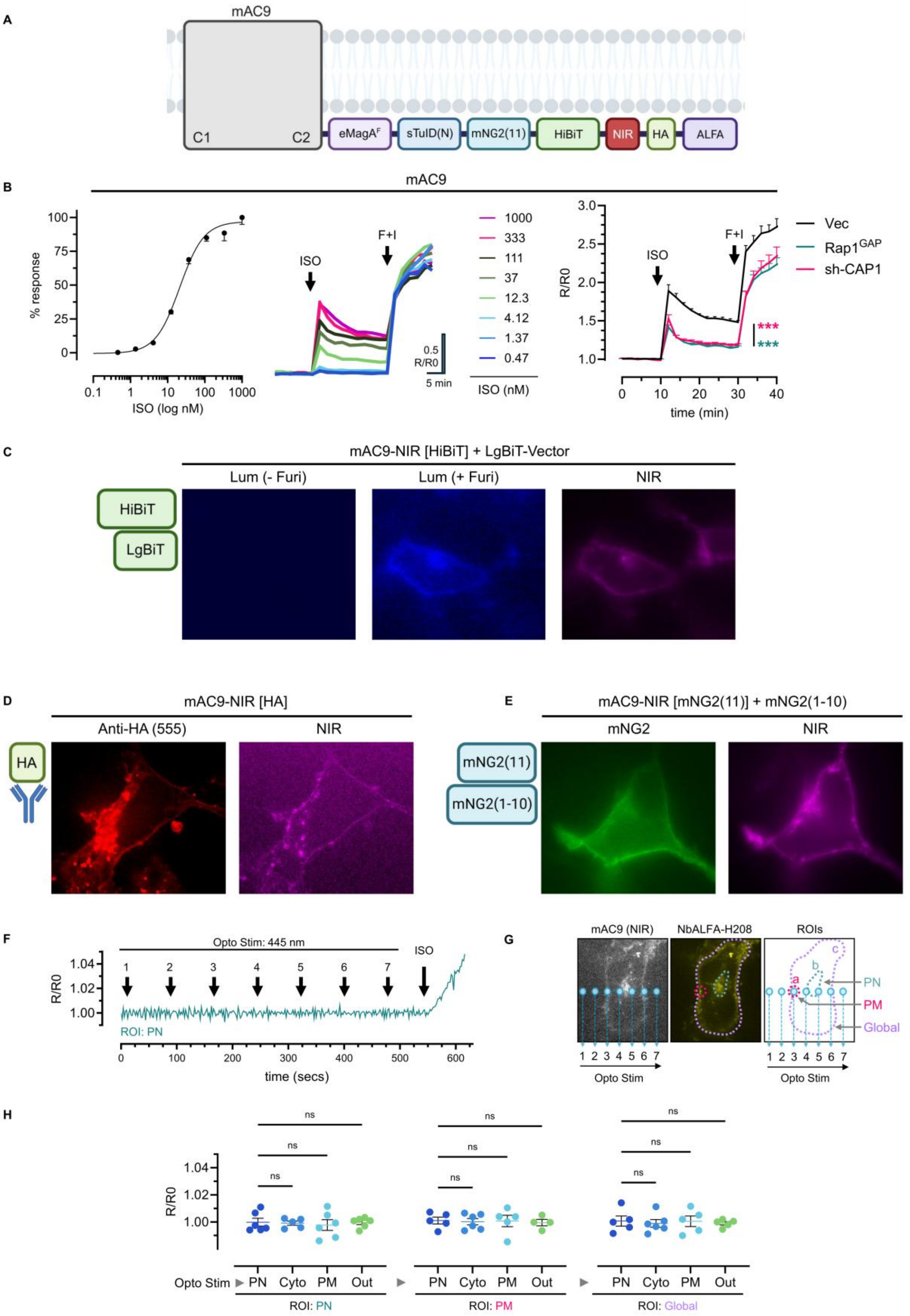
A
mAC9
C1
C2
eMagA^F
sTulD(N)
mNG2(11)
HiBiT
NIR
HA
ALFA
B
mAC9
% response
ISO (log nM)
ISO
F+I
1000
333
111
37
12.3
4.12
1.37
0.47
ISO (nM)
0.5 R/R0
5 min
R/R0
time (min)
Vec
Rap1^GAP
sh-CAP1
***
***
C
mAC9-NIR [HiBiT] + LgBiT-Vector
Lum (- Furi)
Lum (+ Furi)
NIR
HiBiT
LgBiT
D
mAC9-NIR [HA]
Anti-HA (555)
NIR
HA
E
mAC9-NIR [mNG2(11)] + mNG2(1-10)
mNG2
NIR
mNG2(11)
mNG2(1-10)
F
Opto Stim: 445 nm
ISO
R/R0
ROI: PN
time (secs)
G
mAC9 (NIR)
NbALFA-H208
ROIs
PN
PM
Global
Opto Stim
H
ns
R/R0
Opto Stim
PN
Cyto
PM
Out
ROI: PN
ROI: PM
ROI: Global

**Fig. S3** (A–H)**. Design and functional validation of the multi–tagged AC9 (mAC9) construct.** (**A**) Schematic representation of the engineered mAC9 construct. mAC9 was derived from $AC9^{WT}$ by adding the following modules in–frame: $eMagA^{F}$, split–TurboID N–terminal fragment [sTurboID(N)], mNG2(11), HiBiT, NIR (miRFP670nano3 near–infrared fluorescent protein), HA, and ALFA tags. Each module provides an orthogonal readout for localization, interaction mapping, biotinylation–based proximity labeling, complementation assays, or optical detection. (**B**) Dose–response curve of isoproterenol–evoked cAMP production (ISO; 0.47–1000 nM) in HC–1 cells expressing mAC9, derived from real–time FRET traces using the H188 sensor (left panel). Real–time FRET measurements of cytosolic cAMP using the H188 sensor in HC–1 cells expressing mAC9 exposed to increasing concentrations of ISO (0.47–1000 nM) followed by FK plus IBMX (F+I; FK 20 μM, IBMX 250 μM). SEM is omitted for visual clarity (middle panel). cAMP measurements in HC–1 cells expressing mAC9 together with vector control, Rap1GAP, or sh–CAP1. Traces represent mean normalized FRET ratios ($R/R_o$) in response to isoproterenol (ISO; 30 nM) in HC–1 cells expressing mAC9 together with vector control, Rap1GAP, or sh–CAP1 (right panel). (**C–E**) **Validation of mAC9 domains.** (C) HiBiT. Confocal micrographs of an HC–1 cell coexpressing mAC9 and LgBiT–vector show no luminescence in the absence of furimazine (left), robust blue luminescence (PM and perinuclear/endomembrane) after Furimazine addition (10 μM) (middle), and the corresponding NIR fluorescence in the same cell (right). (D) HA–Tag. Immunofluorescence confirmation of HA–tag accessibility. Confocal micrographs of an HC–1 cell expressing mAC9 and stained with anti–HA–555 (left) show PM and perinuclear/endomembrane labeling, with the corresponding NIR fluorescence in the same cell (right). (E) mNG(11). HC–1 cells cotransfected with mAC9 and mNG2(1–10) show reconstitution of green fluorescence (left), and the same cell displays NIR signal (right), confirming proper folding and surface accessibility of the tag. (**F–H**) **Controls for optogenetic activation of mAC9: requirement of $eMagB^{F}$–$Rap1b^{G12V}$.** (F) Representative real–time cAMP measurement using the NbALFA–H208 FRET sensor in HC–1 cells expressing mAC9 during optogenetic blue–light stimulation (445 nm) in the absence of $eMagB^{F}$–$Rap1b^{G12V}$. No cAMP increase was detected in the perinuclear (PN) ROI at any stimulation point. (G) Fluorescence micrographs of an HC–1 cell coexpressing mAC9 and the NbALFA–H208 sensor, showing NIR fluorescence from mAC9 (left) and YFP fluorescence from NbALFA–H208 (middle). The optogenetic stimulation points (1–7) and the ROIs used for FRET quantification (right; PM, plasma membrane; PN, perinuclear; Cyto, cytosolic) are indicated. (H) Quantification of maximal $\Delta R/R_o$ (%) FRET responses during optogenetic stimulation delivered to the indicated stimulation regions (Opto Stim; PN, Cyto, PM, Out) and measured in the specified acquisition ROIs (PN, PM, Global). "Out" denotes stimulation outside the cell. Statistical comparisons were performed by one–way ANOVA with Dunnett's post hoc test versus vector control; ns, not significant. Scatter plots show individual cells with mean ± SEM. Data are representative of ≥3 independent experiments.

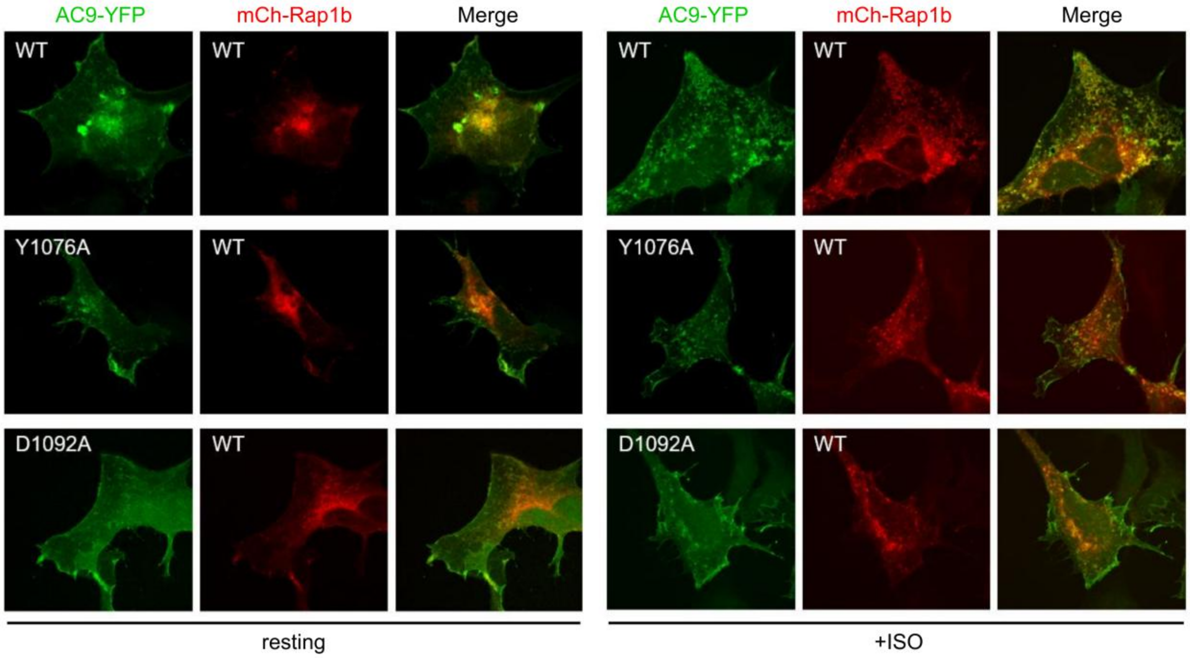


**Fig. S4. Subcellular colocalization of AC9 variants with Rap1b.** Confocal micrographs of HC–1 cells co–expressing AC9–YFP (WT, Y1076A, or D1092A) and mCherry–Rap1b. Micrographs illustrate the subcellular localization and colocalization of Rap1b with $AC9^{WT}$ and AC9 mutants before (resting) and after ISO stimulation (+ISO; isoproterenol 30 nM for 20 min). Cells were fixed, mounted, and imaged by confocal microscopy. Data are representative of n=3 independent experiments.

### I. Adenyl Cyclases and derived constructs

| | *Provided by/Design method* | *Reference/Notes* |
|---|---|---|
| *AC2–HA (rat)* | Dr. Cooper (U. of Cambridge) | |
| *AC2–YFP (bovine)* | Dr. Korkhov (PSI, Zurich, Switzerland) | |
| *AC3 (rat)* | Dr. Dessauer (UTHealth Houston) | |
| *AC5 (canine)* | Dr. Dessauer (UTHealth Houston) | |
| *Flag–AC9 (human)* | Dr. Dessauer (UTHealth Houston) | |
| *AC9–YFP (bovine)* | Dr. Korkhov (PSI, Zurich, Switzerland) | |
| *AC9 WT* | Designed in–house/synthesized by Genescript. | Derived from AC9–YFP (bovine). YFP module removed. |
| *AC9–mCherry* | Designed in–house; synthesized by GenScript. | Derived from AC9–YFP (bovine). YFP module swapped by m–Cherry. |
| *AC2(339)AC9* | Designed in–house; synthesized by GenScript. | AC2(1–P253)–AC9(R339–C–terminus) |
| *AC9(339)AC2* | Designed in–house; synthesized by GenScript. | AC9(1–R339)–AC2(P253–C–terminus) |
| *AC2(764)AC9* | Designed in–house; synthesized by GenScript. | AC2(1–R597)–AC9(R764–C–terminus) |
| *AC9(764)AC2* | Designed in–house; synthesized by GenScript. | AC9(1–R764)–AC2(R597–C–terminus) |
| *AC2(1037)AC9* | Designed in–house; synthesized by GenScript. | AC2(1–P865)–AC9(P1037–C–terminus) |
| *AC9(1037)AC2* | Designed in–house; synthesized by GenScript. | AC9(1–P1037)–AC2(P865–C–terminus) |
| *AC9–YFP–Y1076A* | Designed in–house; synthesized by GenScript. | Point mutations were introduced into bovine AC9–YFP. |
| *AC9–YFP–D1092A* | Designed in–house; synthesized by GenScript. | Point mutations were introduced into bovine AC9–YFP. |
| *Flag–AC9–WT–Y1076A* | Designed in–house; synthesized by GenScript. | Point mutations were introduced into bovine Flag–AC9. |
| *Flag–AC9–WT–D1092A* | Designed in–house; synthesized by GenScript. | Point mutations were introduced into bovine Flag–AC9. |
| *His–C1a* | | See methods |
| *His–C2a* | | See methods |

### II. Rap1 and derived constructs

| | *Provided by/Design method* | *Notes:* |
|---|---|---|
| *HA–Rap1–GAP* | | *Refs. 19 and 20* |
| *HA–Rap1b–WT* | | *Refs. 19 and 20* |
| *HA–Rap1b–G12V* | | *Refs. 19 and 20* |
| *HA–Rap1b–N17* | | *Refs. 19 and 20* |
| *HA–Rap1b–G12V/Q63A* | Designed in–house; synthesized by GenScript. | |

| | | |
|---|---|---|
| *HA–Rap1b–G12V/K73A* | Designed in–house; synthesized by GenScript. | |
| *HA–Rap1b G12V/D69A* | Designed in–house; synthesized by GenScript. | |
| *mCherry–Rap1b WT* | Dr. Philip J. S. Stork (Oregon Health & Science University) | |
| *His–GFP–Rap1b G12V* | Designed in–house; synthesized by GenScript. | See methods |
| *His–Rap1b G12V* | | *Refs. 19 and 20* |
| *Rap1bWT–sTurboID(C)* | Designed in–house; synthesized by GenScript. | |
| *eMagBF–Rap1b G12V* | Designed in–house; synthesized by GenScript. | |
| *HA–Rap1b G12V* | | *Refs 19 and 20* |
| *His–Rap1b–GTPγS* | See methods. | |
| **III. FRET sensors** | | |
| | *Provided by/Design method* | *Notes* |
| *H208* | Dr. Jalink (Netherlands Cancer Institute) | |
| *NbALFA–H208* | Designed in–house; synthesized by GenScript. | Derived from H208. *Ref. 9* |
| *H188* | Dr. Jalink (Netherlands Cancer Institute) | |
| *NLS–H188* | Designed in–house; synthesized by GenScript. | Derived from H188. *Ref. 9* |
| **IV. Other constructs** | | |
| | *Provided by/Design method* | *Notes* |
| *His–GFP* | Designed in–house. | |
| *His–Gαs* | Dr. Korkhov (PSI, Zurich, Switzerland) | |
| *mSA2–EYFP* | Dr.Yubin Zhou's (Texas A&M Health). | *Ref. 34* |
| *Rab5–mNeonGreen* | | Allele Biotech (USA). |
| *sh–CAP1* | | *Refs. 19 and 20* |
| *CAP1* | | *Refs. 19 and 20* |

**Table S1.**
Summary of engineered constructs and modifications.
